# Efficient generation of hematopoietic progenitor cells from human pluripotent stem cells by robotic automation

**DOI:** 10.64898/2026.04.07.716925

**Authors:** Kenjiro Adachi, Noriyasu Okazaki, Ayato Sugiyama, Yota Goto, Fumika Shimamura, Yuuwa Takahashi, Masayuki Ito, Atsushi Inoue, Hideto Yamaguchi

## Abstract

Differentiation of human pluripotent stem cells (hPSCs) into therapeutic cell types requires stringent control of developmental signals, but conventional manual operations introduce substantial intra- and inter-operator variability. While automation can reduce such variability, fixed systems often lack compatibility with the dynamic, stage-specific, and format-diverse manipulations required for hPSC culture and differentiation. Here, we establish a flexible robotic platform to standardize a stroma- and xeno-free embryoid body (EB)-based hematopoietic progenitor cell (HPC) differentiation process and integrate machine learning (ML) to optimize key process parameters including the concentrations of multiple signaling molecules. Despite intrinsic biological stochasticity and technical complexity associated with EB formation, the robotic platform substantially reduced experimental variation. ML-guided, unbiased exploration not only identified combinations of signaling inputs that markedly improved the efficiency and reproducibility of HPC differentiation and subsequent natural killer cell generation, but also illuminated key signaling logic underlying early human hematopoietic development. Together, these results demonstrate that data-driven robotic automation can uncover optimal culture conditions that are difficult to identify through conventional human-driven experimentation.

## Introduction

Allogeneic cell therapies using human pluripotent stem cells (hPSCs) hold great promise for both cell replacement and immunotherapeutic approaches for cancer and autoimmune diseases, offering innovative solutions to previously untreatable conditions. These therapies leverage the unique ability of hPSCs to differentiate into any cell type, making them well suited for restoring function in damaged tissues. hPSCs are also highly amenable to genetic engineering, enabling the generation of allogeneic cells with reduced immunogenicity and enhanced functionality, persistence, or tumor specificity. Preparing a large number of engineered immune cells as off-the-shelf therapeutics could make cancer treatments more accessible and affordable. Numerous clinical trials evaluating hPSC-derived products are currently ongoing^1^, targeting a wide range of therapeutic areas including neurodegenerative diseases, cardiovascular disorders, ocular diseases, and cancers. While most of these trials remain in early phases and primarily assess safety, tolerability, and preliminary efficacy, several studies have reported encouraging evidence of engraftment and functional improvement.

Despite these advances, significant challenges remain in translating hPSC-derived therapies to the clinic, particularly those related to the biological variability and complexity of manufacturing processes. The development of cell therapies involves several phases, from discovery through preclinical and clinical research to process development and manufacturing, each of which can introduce biological, technical, or procedural changes that impact the consistency and quality of the final product. The self-renewal and differentiation of hPSCs are highly sensitive to both chemical and physical cues and typically rely on a series of non-standardized manual operations, often leading to inconsistent outcomes. The introduction of new operators during development, combined with insufficient training, can further increase variability in culture, differentiation, and expansion techniques, compromising reproducibility. Suboptimal differentiation protocols can also lead to heterogeneous cell populations and variable functional outcomes, thereby undermining the consistency and quality of final products. As products progress from research to clinical development, materials, manufacturing processes, and analytical methods often require modification to meet evolving regulatory and manufacturing requirements, potentially impacting product quality and consistency.

To address these complex sources of variability and the biological complexity inherent to living therapeutics, we explore the integration of robotic automation and artificial intelligence (AI)-driven process optimization into cell therapy manufacturing. Robotic platforms can standardize critical operations such as cell seeding, media exchanges, and differentiation, thereby minimizing operator-dependent variability and enhancing reproducibility. The establishment of a robust differentiation process from early developmental stages is crucial, as it critically affects cellular functionality, differentiation efficiency, and the presence of off-target lineages, potentially compromising both efficacy and safety if not properly controlled. Moreover, once the program advances to the clinical phase, modifications to the differentiation process are difficult and require extensive justification. Early implementation of a reproducible process is also essential for scaling up to clinical-grade manufacturing, ensuring consistent product quality and generating reliable data for regulatory submissions. However, optimization of multiple process parameters typically requires a large number of experiments and considerable time. This is mainly due to the high dimensionality of the search space, complex and often non-linear interactions among variables, and the risk of getting trapped in local maxima rather than identifying the true global optima. Machine learning (ML)-based systems, capable of analyzing high-dimensional datasets and extracting predictive insights, offer a robust framework for addressing these challenges and accelerating process development. Black-box optimization (BBO) has become increasingly important for efficiently searching optimal process conditions without requiring explicit knowledge of the underlying model. BBO aims to identify the optimum of an unknown function describing the relationship between process conditions and outcomes by iteratively exploring and exploiting information obtained from previous experiments. Among these approaches, Bayesian optimization (BO) is a powerful method that builds a probabilistic surrogate model and iteratively updates it as new data are collected. By balancing exploration and exploitation through an acquisition function, BO efficiently identifies high-performing regions of an expensive black-box objective and can approach the global optimum under appropriate modeling assumptions. Such data-driven optimization frameworks are particularly valuable in stem cell biology, where subtle variations in culture parameters can profoundly influence cell fate determination and functional maturation. The synergistic application of automation and AI has the potential not only to improve reproducibility and scalability but also to accelerate the development of streamlined manufacturing processes.

In this study, we combined robotic automation with an AI-based optimization framework to maximize the differentiation efficiency of hPSCs into hematopoietic progenitor cells (HPCs) via embryoid body (EB) formation. This approach enabled the identification of new culture conditions that reproducibly and efficiently generate HPCs from multiple human induced pluripotent stem cell (hiPSC) lines through unbiased exploration of a large experimental design space. These HPCs were functionally validated by their efficient differentiation into natural killer (NK) cells exhibiting antitumor cytotoxicity under stroma-and xeno-free conditions.

## Results

### Synergistic integration of robotic automation and BBO for multiparametric process optimization

hPSC-derived NK cells represent a promising platform for allogeneic off-the-shelf immunotherapies. However, differentiation protocols frequently rely on co-culture with murine stromal cells to support hematopoietic and NK cell development, introducing xenogeneic components and complicating clinical translation. Moreover, the clinical-scale manufacturing of NK cells is constrained by the low efficiency of HPC differentiation from hPSCs and the downstream NK cell production often shows substantial batch-to-batch variability. We have developed a xeno-free NK cell differentiation protocol (details are described in a manuscript in preparation) based on the published stroma-free method^2^. This protocol (hereafter called “standard” method) involved the formation of EBs in the presence of BMP4, VEGF, and SCF to differentiate hPSCs into HPCs, followed by culturing on DLL4- and RetroNectin-coated plates in the presence of cytokines and chemical compounds that promote progenitor cell proliferation and NK cell differentiation. Although this protocol enables the generation of NK cells with typical NK cell-related functions, such as natural cytotoxicity against tumor cells and proinflammatory cytokine production in response to tumor cells, substantial variability in NK cell yield persists. This variability is observed both across independent experiments and among different hPSC lines, possibly due to a combination of human-induced differences in culture and differentiation techniques, intrinsic biological stochasticity associated with EB-mediated differentiation including heterogeneity in the local microenvironment driven by direct cell-cell interactions and paracrine signaling among distinct cell populations, as well as suboptimal aspects of the differentiation protocol.

To develop a more robust and efficient NK cell differentiation protocol suitable for clinical translation, we attempted to standardize the EB formation process and improve the consistency by applying a robotic automation system. Our robotic platform, Maholo^3^, was designed to automate a variety of cell culture processes using a programmable robot with high precision and flexibility. It consists of a dual-arm robot that can imitate natural human motions and operate a number of common laboratory instruments and equipment such as incubators, centrifuges, microscopes, cell counters, refrigerators, dry baths, micropipettes, and aspirators (Fig. 1a). This flexible robotic platform enables the automation of various differentiation steps with minimal modifications to established methods, particularly during the early stages of drug development. Such automation reduces variability, promotes process standardization, improves data reliability, and enables the consistent execution of complex procedures with minimal human error, facilitating more accurate and rapid process optimization.

**Fig. 1.**
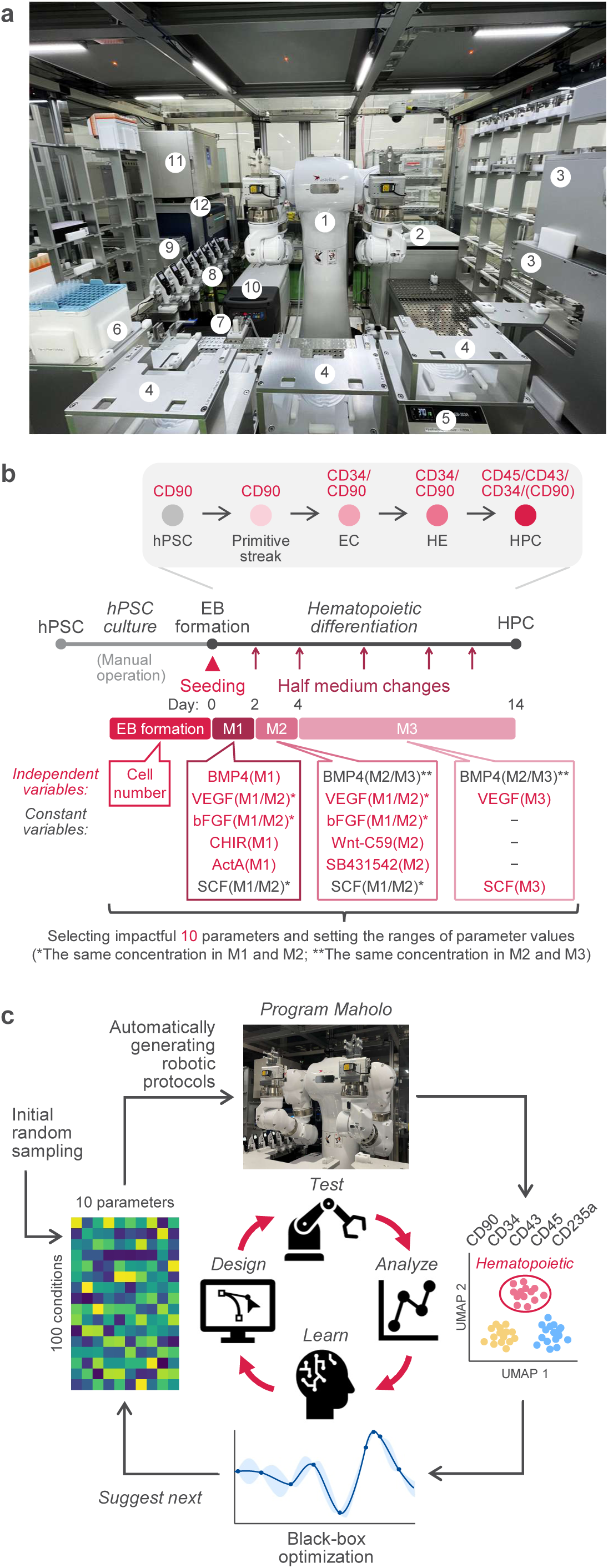
The Maholo robotic system for process optimization. **a**, An overview of the robotic platform. The robot and peripheral equipment are enclosed in a clean-air environment with HEPA-filtered airflow directed over the incubator and open handling area. The central dual-arm robot (1) can access all peripheral devices, equipment and materials, including the centrifuge (2), microscopes (3), operation racks (4), block bath (5), tip racks (6), aspirator (7), electronic pipettes (8), conveyor to the incubator (9), freezer (10), cool incubator (11), and cell counter (12). **b**, Schematic illustrating the stepwise differentiation of hPSCs into HPCs. Cell seeding into microwell plates, EB formation, and sequential media exchanges were automated using the robotic platform. To identify optimal combinations of signaling inputs for efficient HPC differentiation, the following 10 parameters were selected as independent variables: the number of cells in each EB, the concentrations of BMP4, CHIR, and ActA in mesoderm induction medium (M1), those of Wnt-C59 and SB431542 in endothelial differentiation medium (M2), those of VEGF and bFGF in M1/M2, and those of VEGF and SCF in hematopoietic differentiation medium (M3). We held the concentration of SCF in M1/M2 and that of BMP4 in M2/M3 constant across all conditions. **c**, Schematic of the optimization of process parameters through statistical ML.

We translated the manually operated EB formation procedure into an automated workflow by programming the robot to perform iPSC seeding into microwell plates and a series of media exchanges to drive stepwise differentiation into posterior primitive streak/lateral plate mesoderm, endothelial cells, hemogenic endothelium (HE), and HPCs (Fig. 1b, Extended Data Fig. 1a-d, and Supplementary Videos 1-3). We sought to modulate the local microenvironment by temporally controlled exposure to external cues. In addition to BMP, VEGF, and SCF signaling, the WNT, Activin/Nodal, and FGF signaling pathways also play key roles in the formation of the primitive streak and the induction of mesoderm^4^ as well as in the progressive development of the endothelial and hematopoietic lineages^5^. Thus, we treated EBs with combinations of cytokines and chemical compounds that modulate these pathways and examined the expression of surface markers characteristic of hPSCs and primitive streak/early mesoderm (CD90), endothelial cells including HE (CD34/CD90), HPCs (CD34/CD43/CD45), as well as CD235a, which is expressed in primitive hematopoietic progenitors^6^. Preliminary experiments conducted manually suggest that activation of WNT, Activin/Nodal, and FGF signaling pathways in the presence of BMP4, VEGF, and SCF can promote the emergence of CD43^+^ hematopoietic cells. However, these experiments did not identify optimal combinations and concentrations of these signaling molecules that would increase the yield of HPCs competent for NK cell differentiation.

Guided by the preliminary results, we decided to optimize the concentrations of signaling molecules and the number of hPSCs for EB formation using a ML approach based on BO, which was previously applied to promote hPSC differentiation into retinal pigment epithelium^7^. We first defined the combinations and timings of signaling pathways activated or inhibited during stepwise differentiation of hPSCs into HPCs. Feeder-free hiPSCs were dissociated into single cells, reaggregated in microwell plates to facilitate formation of EBs with uniform size, and then cultured in mesoderm induction medium (M1) containing BMP4, VEGF, bFGF, CHIR99021 (CHIR; WNT/β-catenin activator), Activin A (ActA), and SCF for two days. At day 2, half of the medium was replaced with fresh endothelial differentiation medium (M2) containing BMP4, VEGF, bFGF, Wnt-C59 (PORCN/WNT inhibitor), SB431542 (Activin/Nodal/TGF-β inhibitor), and SCF. On days 4, 7, 10, and 12, half of the medium was replaced with fresh hematopoietic differentiation medium (M3) containing BMP4, VEGF, and SCF. We set the 10 most critical variables as optimization parameters, thereby constraining the search space and enabling efficient exploration with minimal iterations.

Our automated workflow enabled parallel testing of 100 or more conditions (Extended Data Fig. 1e). The parameter values for the 100 experimental conditions in the first iteration were selected by random sampling (Fig. 1c). Robotic protocols were automatically generated based on these parameter values, with the 100 conditions distributed across five plates. On day 14, EBs generated by the semi-automated system were manually harvested, dissociated into single cells, and analyzed for the expression of CD90, CD34, CD43, CD45, and CD235a by flow cytometry. We evaluated the hematopoietic differentiation efficiency for each sample by quantifying the percentage of CD43^+^ cells, which was then used as the objective variable to be maximized. The surrogate model was trained with the data from the first iteration and the acquisition function was used to propose the most promising set of parameters for the next iteration. Through repeated cycles of design, testing, and learning, the model is iteratively updated with new data and progressively converges on the optimal parameter combination.

### Accelerated development of an EB-based protocol for efficient hematopoietic differentiation of hiPSCs

We conducted three optimization iterations, evaluating 246 unique conditions across 300 samples. The optimization process can become less efficient and take longer to converge when the experimental noise is high, because the model may overexplore noisy regions, misinterpret random fluctuations as meaningful patterns, or become trapped around false optima. We assessed the variability of cell populations generated under identical conditions. Despite the varying proportions of different cell populations, we observed excellent consistency in cell population distributions both within and across experiments (Extended Data Fig. 2a–c). As a result, each observation provided maximal information for modeling the objective function.

Next, we performed automated analysis of large, multi-dimensional flow cytometry datasets and identified three major cell populations: endothelial cells (CD34^+^/CD90^+^), hematopoietic cells (CD43^+^), and other populations (CD34^−^/CD43^−^) (Fig. 2a and Supplementary Table 1). The proportion of CD43^+^ cells (“a hematopoietic score”) was calculated for each sample and used as an objective metric for optimization. In the first iteration with random sampling, we found several conditions under which a surprisingly large proportion of cells were differentiated into hematopoietic cells, although the majority of conditions failed to improve hematopoietic differentiation compared with the standard condition (Fig. 2b). We tested 78 and 72 new conditions proposed by BO in the second and third iterations, respectively, and found that most of the conditions proposed by the model gave near-optimum outcomes after three iterations of optimization (Fig. 2b and Supplementary Tables 2 and 3), indicating that the algorithm had identified the promising region and that further sampling did not appear to improve outcomes significantly. Strikingly, microscopic observations revealed that the EBs gradually became disorganized and round hematopoietic-like cells emerged and spread into the surrounding area when a large proportion of cells were differentiated into CD43^+^ cells (Fig. 2c). In summary, our robotic system provides a highly reproducible EB-based differentiation platform that, together with BO, enables the rapid identification of multiple conditions yielding hematopoietic cells with nearly 90% efficiency.

**Fig. 2.**
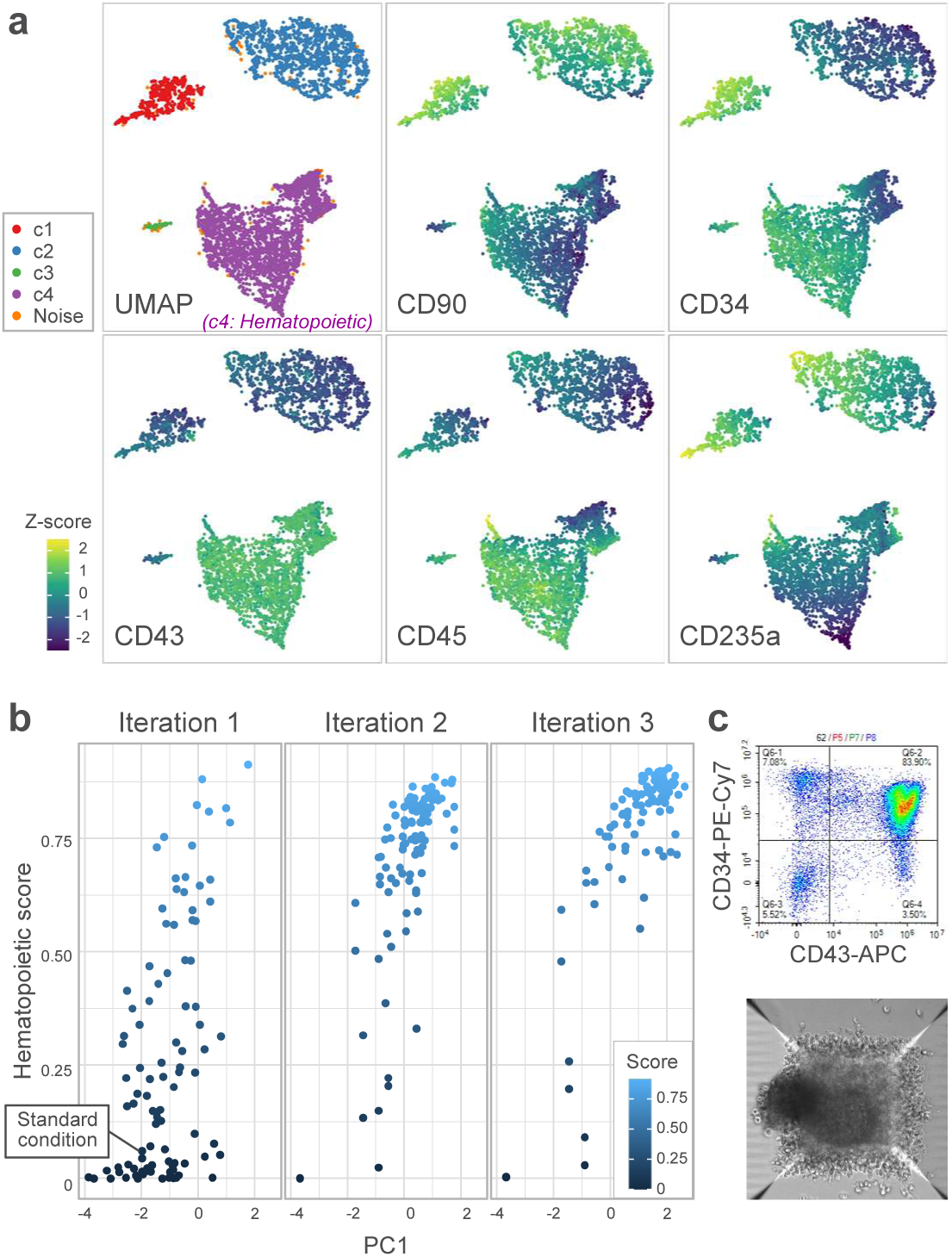
|Rapid optimization of process parameters for efficient generation of hematopoietic cells from hiPSCs. **a**, Cell populations visualized by UMAP. To process flow cytometry data, debris, doublets, dead cells, and lineage-positive cells were excluded and 1,000 cells were randomly selected from each sample. CD90, CD34, CD43, CD45, and CD235a were used to perform dimensionality reduction with UMAP followed by clustering with HDBSCAN^19^. The colored plots show Z-score-normalized expression levels of the indicated markers. The broad cluster with CD43 expression (c4) was annotated as hematopoietic cells and a hematopoietic score for each sample was calculated as the proportion of hematopoietic cells multiplied by the proportion of viable cells. Noise, data points that are not assigned to any cluster by HDBSCAN. **b**, Scatter plot showing hematopoietic scores for each sample across three iterations. Partial Least Squares (PLS) regression was used to model the relationship between multiple process parameters and hematopoietic scores. PLS identifies latent variables that represent weighted combinations of process parameters showing the strongest covariance with the hematopoietic scores, with the first component (PC1) capturing the parameter pattern most strongly associated with the scores. Only new conditions proposed by BO in each iteration are plotted. **c**, A representative flow cytometry plot (top) and phase-contrast microscopy image (bottom) of samples at day 14 showing high hematopoietic scores.

### Two divergent pathways of hematopoietic differentiation revealed by unbiased flow cytometry analysis

We next took an unbiased, data-driven approach to uncover patterns and insights from the complex multi-dimensional flow cytometry data from 300 samples. We used FlowSOM^8^, an unsupervised clustering algorithm based on a self-organizing map (SOM), to detect subpopulations present in the EBs on day 14. By training the SOM with the cell surface marker profiles (CD90, CD34, CD43, CD45, and CD235a), the cells were clustered into 49 nodes, which were further grouped into 15 metaclusters using consensus clustering (Fig. 3a). These metaclusters represent different cell types or cellular states. Metaclusters 4, 5, and 6 are characterized by co-expression of CD34, CD43, and CD45, and are thus considered putative HPCs (Fig. 3b-d). Metaclusters 7, 9, 11, and 12 are negative for one or more of CD34, CD43, or CD45, likely representing more differentiated hematopoietic cells (referred to hereafter as “Hema”). Metaclusters 3 and 15 are considered erythroid cells based on their high expression of CD235a. Metaclusters 1 and 2 exhibit high levels of the mesenchymal marker CD90 and the endothelial marker CD34, suggesting their endothelial identity. The remaining metaclusters (8, 10, 13, and 14) lack CD34 as well as the hematopoietic markers CD43 and CD45, apparently corresponding to other mesodermal or non-mesodermal cell types.

**Fig. 3.**
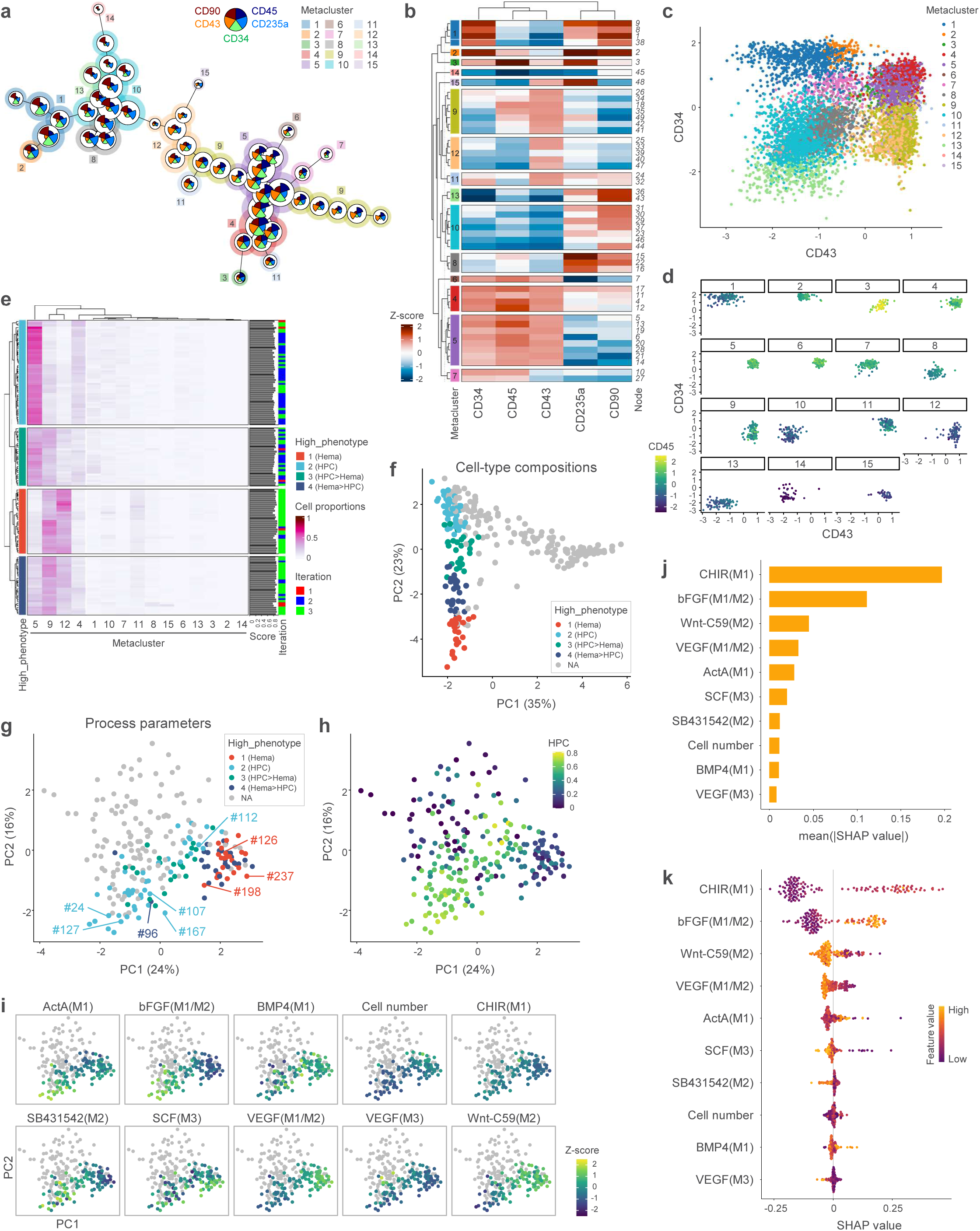
Two divergent hematopoietic differentiation programs driven by distinct combinations of signaling inputs. **a**, FlowSOM minimum spanning tree with 49 nodes and 15 metaclusters. Each node is connected with its most similar neighbor. **b**, Heatmap showing median expression levels of the markers for each node. **c**, Dot plot of CD43 and CD34 expression in 10,000 randomly sampled cells. Different colors correspond to different metaclusters. **d**, Dot plots showing CD43, CD34, and CD45 expression in the cells of each cluster. One hundred cells were randomly sampled from each cluster for visualization. **e**, Heatmap showing compositions of metaclusters for each of the 137 samples with the hematopoietic score ≥ 0.75. Four phenotypic groups were identified by partitioning around medoids (PAM) clustering based on Jensen-Shannon distance (JSD) among samples. **f**, PCA plot based on compositions of metaclusters for all 300 samples. The samples with the score < 0.75 (NA for high-score phenotype groups) are shown in gray. The percentages of variance explained are shown in brackets. **g**, PCA plot based on process parameters. The samples with the hematopoietic score < 0.75 are shown in gray. Parameter IDs are shown for the selected conditions. **h**, PCA plot of process parameters with colors representing proportions of HPC (metaclusters 4/5/6). **i**, PCA plot of process parameters with colors representing Z-score-normalized parameter values. The samples with the hematopoietic score < 0.75 are shown in gray. **j**, Mean absolute SHAP values indicating average impact of each feature on the prediction of HPC fate. **k**, SHAP beeswarm plot showing the directional impact of individual features on the prediction of HPC fate across samples. Positive SHAP values push the prediction toward HPC fate, while negative values push the prediction toward the other.

We clustered all 300 samples based on their cell-type composition profiles and obtained three phenotypic groups, each characterized by a high proportion of either HPCs, Hema, or other cell types (Extended Data Fig. 3a and Supplementary Table 2). Samples containing erythroid cells (i.e., Parameter ID #96 and #189) were also enriched in Hema and therefore classified into the Hema group. To further investigate the samples exhibiting high hematopoietic differentiation efficiency, 137 samples with the hematopoietic score ≥ 0.75 were selected and subsequently clustered into four phenotypic groups (Fig. 3e). The first and second groups consist predominantly of samples enriched in Hema (metaclusters 7/9/11/12) and HPCs (metaclusters 4/5/6), respectively. In contrast, the remaining two groups represent mixed populations containing both Hema and HPCs. These results suggest that hiPSCs undergo hematopoietic differentiation through two divergent pathways, influenced by the concentrations of signaling molecules and the number of cells in each EB. The principal component analysis (PCA) of cell-type composition profiles showed that samples with high and low hematopoietic differentiation efficiency are clearly separated along the first principal component, whereas samples enriched in HPCs and Hema are separated along the second principal component (Fig. 3f and Extended Data Fig. 3b).

We next sought to identify key factors determining the two divergent hematopoietic fates. The PCA of process parameters revealed that samples enriched in HPCs and Hema (groups 2 and 1 of the hematopoietic score ≥ 0.75, respectively) as well as those with low hematopoietic differentiation efficiency (the hematopoietic score < 0.75) cluster separately (Fig. 3g,h and Extended Data Fig. 3c), indicating that the process parameter sets are similar within each group but differ substantially among groups. The low hematopoietic score group is characterized by relatively high cell numbers and high levels of VEGF in the hematopoietic differentiation medium (M3) (Fig. 3h and Extended Data Fig. 3d–i). The HPC group is associated with low cell numbers, low levels of VEGF in M3, high levels of bFGF in the mesoderm induction medium (M1) and endothelial differentiation medium (M2), and high levels of SB431542 in M2, while the Hema group is associated with low cell numbers, low levels of VEGF in M3, high levels of VEGF in M1, and high levels of Wnt-C59 in M2.

We further investigated the parameters determining HPC fate relative to Hema using a ML approach. For this, a random forest algorithm was chosen for its high interpretability, enabling evaluation of parameter contributions through feature importance and SHapley Additive exPlanations (SHAP) analysis^9^. We first selected the 137 samples with high hematopoietic score and trained a random forest classifier to distinguish HPC (high-score phenotype group 2) from other groups based on the parameter values. We performed a 10-fold cross-validation to evaluate model performance and identify the optimal hyperparameter combination. The cross-validation achieved an accuracy of 0.906, indicating that the model was well suited to distinguishing HPC from other phenotypes. After hyperparameter tuning, the final random forest model was trained on the entire dataset using the optimal parameters. The importance of each parameter in the final model was estimated using built-in feature importance metrics. CHIR in M1 was identified as the most influential parameter for HPC, followed by bFGF in M1/M2 and Wnt-C59 in M2 (Extended Data Fig. 4a). We next assessed the contribution of each parameter to the model’s predictions for individual samples using SHAP values. SHAP values show both the magnitude and direction of each feature’s effect on a given prediction. The top three parameters with the highest mean absolute SHAP values, which represent the global importance, were CHIR in M1, bFGF in M1/M2, and Wnt-C59 in M2 (Fig. 3j), consistent with the feature importance results. At the local level, intermediate concentrations (2–3 µM) of CHIR in M1 increase the predicted probability of HPC fate (high positive SHAP values), whereas low levels of CHIR decrease it (high negative SHAP values) (Fig. 3k and Extended Data Fig. 4b,c). In addition, higher bFGF in M1/M2, as well as lower Wnt-C59 in M2, enhance the prediction of HPC fate. These results highlight the critical role of key signaling molecules in determining HPC fate. Considering the presence of CD235a^+^ cells in Hema but not in HPC conditions, and in light of previous reports linking WNT and Activin/Nodal signaling to definitive and primitive hematopoiesis, respectively^6,10^, HPCs and Hema might arise from the corresponding lineages.

### Highly efficient generation of NK cells from CD34^+^/CD43^+^ HPCs

As a functional validation of HPCs, we first induced their differentiation into NK cells under stroma- and xeno-free conditions. EB-derived cells were seeded onto DLL4- and RetroNectin-coated plates and cultured for 3–4 weeks in the presence of SCF, IL-7, IL-15, FLT3L, and UM171 (Fig. 4a). HPCs generated using our protocol were efficiently and reproducibly differentiated into CD56^+^/CD45^+^/CD3^−^ NK cells (Fig. 4b-e and Extended Data Fig. 5a). Conditions that generated a larger number of HPCs (metaclusters 4/5/6) yielded more NK cells, whereas conditions that produced fewer HPCs resulted in a smaller number of NK cells. Interestingly, conditions that predominantly produced Hema with or without erythroid cells (Parameter ID #96, 126, and 186) failed to generate NK cells, underscoring the functional divergence of HPCs and Hema. In one of the optimized conditions (Parameter ID #24), 17–28 million NK cells were obtained per 15,000 hiPSCs after 2–3 weeks of NK cell differentiation, which is approximately 50–100-fold more efficient compared with the published stroma-free protocols (approximately 3 million NK cells per 300,000 hiPSCs after 4 weeks^11^ or 2–20 million NK cells per 768,000 hiPSCs after 3-5 weeks^2^).

**Fig. 4.**
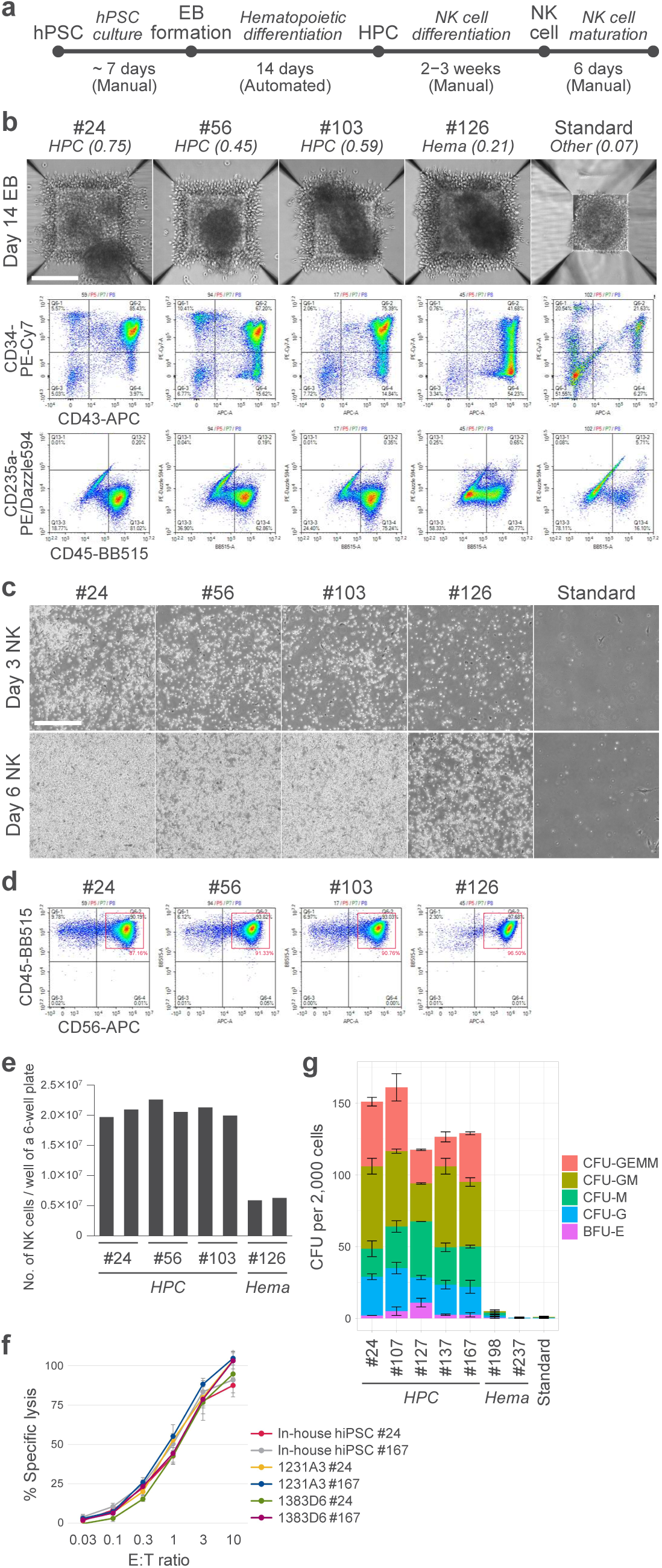
Robust and efficient NK cell differentiation from hiPSC-derived HPCs. **a**, Stroma- and xeno-free NK cell differentiation from hPSCs. **b**, Representative phase-contrast microscopy images (top) and flow cytometry plots (middle and bottom) of day 14 EBs. Phenotype groups (1: HPC, 2: other, 3: Hema) and proportions of HPC are shown for each parameter ID. Only the first replicate is shown for each condition. Scale bar, 200 µm. **c**, Representative images of cells three and six days after seeding under NK cell differentiation conditions. Scale bar, 500 µm. **d**, Representative flow cytometry plots of day 14 NK cells. The standard condition yielded too few cells for analysis. Only the first replicate is shown for each condition. **e**, The number of NK cells obtained from a single well of a six-well plate. Values were calculated as the number of viable cells multiplied by the proportion of CD56^+^/CD45^+^ cells. EB formation and NK differentiation were performed in duplicate for each condition. **f**, Natural cytotoxic activity of NK cells against K562 cells. Mature NK cells derived from three different hiPSC lines under two different HPC conditions were cocultured with K562-HiBiT cells at various effector-to-target (E:T) ratios for 4 hours. HiBiT-tagged proteins released from lysed target cells were measured and presented as the percentage of specific lysis compared with maximum lysis in digitonin. Data represent mean ± SD of triplicates. **g**, CFU assay of cells derived from day 14 EBs. Data represent mean ± SD of duplicates.

We assessed the robustness and reproducibility of our method using three additional hiPSC lines. These hiPSCs were induced to differentiate into HPCs through EB formation and subsequently into NK cells. Two out of three lines were efficiently differentiated into HPCs and NK cells under the optimized conditions for the initial hiPSC line (Extended Data Fig. 5b,c), thereby reinforcing the robustness of our method. These lines also responded similarly to the complex signal inputs as the initial line, committing to either HPC or Hema/erythroid cell fates. One hiPSC line 409B2 was refractory to the hematopoietic differentiation (Extended Data Fig. 5b–e), consistent with the previous studies^12^.

To assess the functionality of hiPSC-derived NK cells, we first cultured the NK cells on plates coated with anti-NKp30 and anti-DNAM-1 antibodies and RetroNectin in the presence of IL-15, IL-18, and IL-21, followed by suspension culture in the presence of IL-15 and IL-18 to promote NK cell maturation. The mature NK cells expressed activating NK cell receptors such as NKG2D and DNAM-1 (Extended Data Fig. 6a). These cells contained a small percentage of CD56^+^/CD16^+^ population, a feature commonly observed in hPSC-derived NK cells^2, 11, 13^. We then performed cytotoxicity assays using K562 leukemia cells to evaluate their natural cytotoxicity. NK cells derived from three different hiPSC lines under two different HPC conditions were all capable of killing K562 cells consistently (Fig. 4f). We also confirmed that NK cells secreted IFN-γ and TNF-α upon the stimulation with K562 cells (Extended Data Fig. 6b), indicating that NK cells derived from the EBs differentiated with the optimized protocol are functional.

We next performed colony-forming unit (CFU) assays to assess the ability of HPCs to differentiate into myeloid and erythroid lineages. Seeding 2,000 unsorted cells resulted in 100–160 colonies for each of the five different HPC conditions, 15-30% of which were mixed colonies (CFU-GEMM) (Fig. 4g and Extended Data Fig. 6c). In contrast, the cells differentiated under the Hema conditions failed to form hematopoietic colonies. Taken together, these results indicate that hiPSCs were efficiently differentiated into functional HPCs.

### Emergence of HPCs through self-organized symmetry breaking

To dissect the mechanism governing the emergence of HPCs from hPSCs, we first examined the contribution of key signaling pathways by omitting individual cytokines and chemical compounds under the two different HPC conditions: one with smaller EBs (500 cells per EB for Parameter ID #24) and another with larger EBs (1,600 cells per EB for Parameter ID #112; Extended Data Fig. 7a). Flow cytometry analysis revealed that BMP4 in M1, bFGF in M1/M2, and VEGF in M2 are critical for HPC induction in both conditions, and removal of SCF from M3 had a moderate effect (Fig. 5a and Extended Data Fig. 7a,b). Removing CHIR from M1 greatly reduced CD43^+^/CD45^+^ hematopoietic cells, while the simultaneous removal of CHIR and Wnt-C59 only moderately affected the induction of hematopoietic cells although CD34^+^ HPCs were significantly reduced, highlighting the importance of endogenous canonical and/or noncanonical WNT ligands. Consistent with the requirement for intermediate levels of CHIR in the HPC group (Fig. 3k and Extended Data Fig. 3e,h), both an increase and a decrease in CHIR concentration under one of the two HPC conditions (Parameter ID #24), as well as a decrease in CHIR concentration under the other condition (Parameter ID #112), resulted in a significant reduction of HPCs. Although the removal of ActA from M1 did not seem to affect the percentage of HPCs detected by flow cytometry, most of the EBs formed in the absence of ActA were small, with a few EBs developed to those containing many hematopoietic-like cells, and the total number of HPCs was markedly reduced in one HPC condition (Parameter ID #24; Fig. 5a and Extended Data Fig. 7b,c). The importance of exogenous ActA seems context-dependent, as its removal in another condition (Parameter ID #112) did not affect the percentage of HPCs and even resulted in a significant increase in HPC number. We confirmed that the number of CD34^+^/CD43^+^ HPCs was largely correlated with the capacity to generate NK cells (Extended Data Fig. 7d,e).

**Fig. 5.**
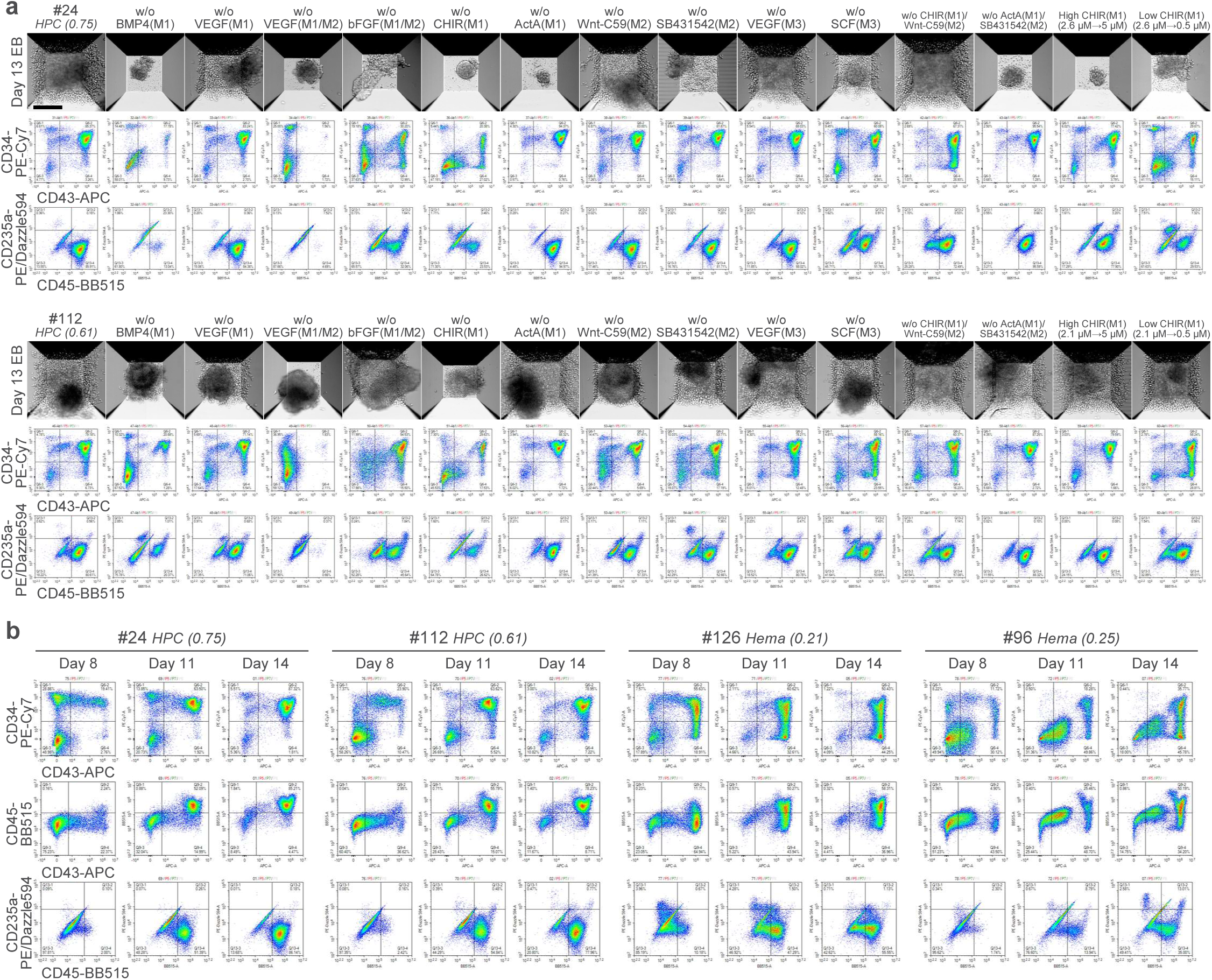
Tightly regulated mechanisms that orchestrate hematopoietic differentiation. **a**, Effects of perturbing key signaling inputs on HPC phenotype and function. Representative NAMC microscopy images (top) and flow cytometry plots (middle and bottom) of day 14 EBs are shown. Scale bar, 200 µm. **b**, Time-course flow cytometry analysis of EBs cultured under the HPC and Hema conditions.

We then sought to elucidate the sequence of events leading to HPC formation. Flow cytometry analysis of EBs at earlier time points suggested that HPC development proceeds through a transition from CD34^+^/CD43^−^ endothelial cells to CD34^+^/CD43^+^ hematopoietic cells around day 8, and subsequently to CD34^+^/CD43^+^/CD45^+^ HPCs by day 11, in a temporally regulated manner under continuous exposure to the same culture medium from days 4 to 14 (Fig. 5b). Under the Hema conditions, however, the CD34^+^/CD43^+^ hematopoietic cells may have emerged earlier than day 8, consistent with the fact that during development, primitive hematopoiesis occurs earlier than definitive hematopoiesis.

We next focused on the morphological changes of EBs during the 14 days of differentiation. The EBs cultured under the HPC conditions showed dynamic morphological changes. They developed into elongated structures around day 6–8 and hematopoietic-like cells emerged shortly thereafter from larger lobes of elongated EBs (Fig. 6a). In contrast, EBs cultured under the Hema conditions were not elongated and irregularly shaped, yet hematopoietic-like cells emerged as early as day 6.

**Fig. 6.**
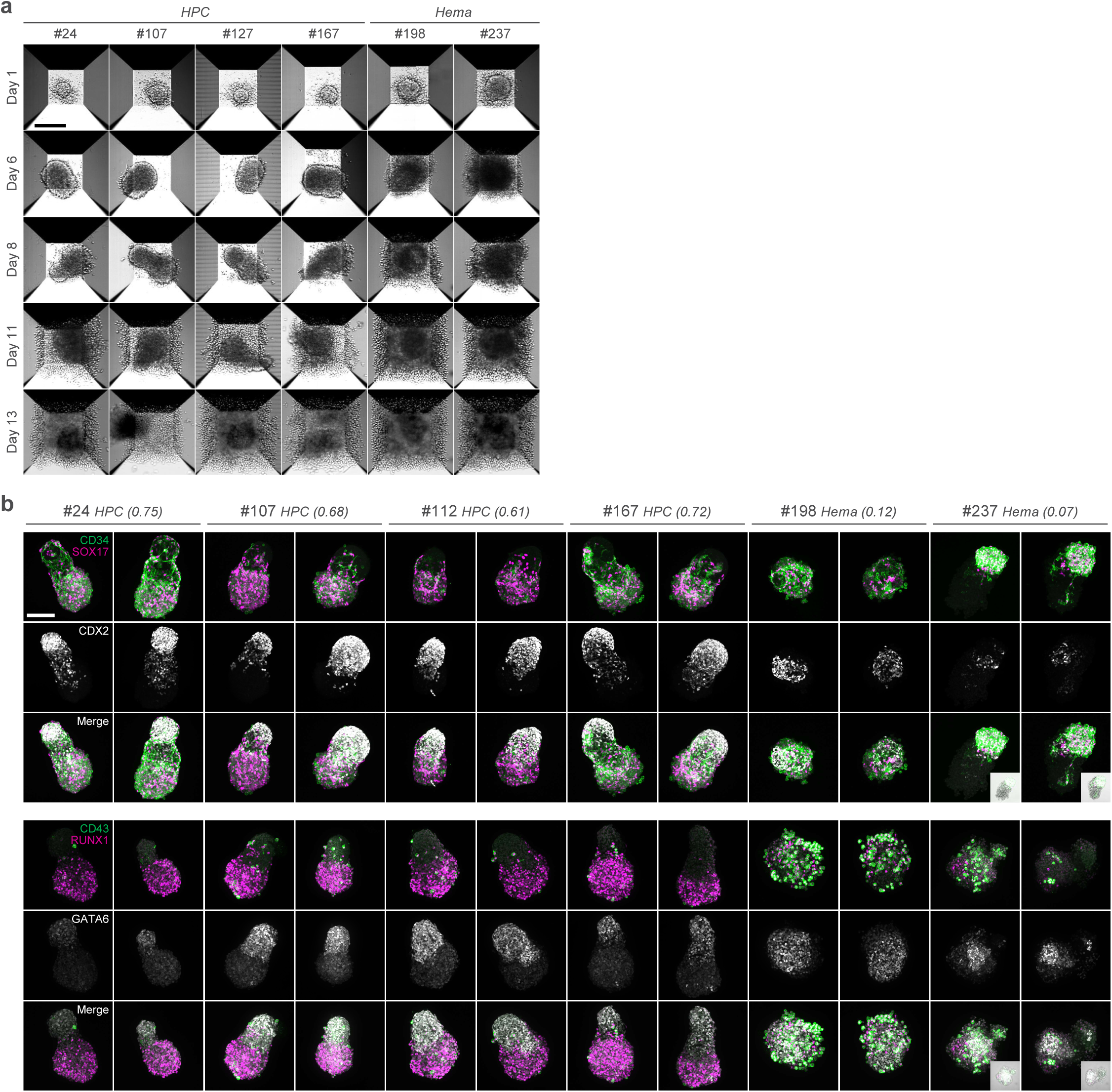
Formation of elongated EBs with polarized expression of key transcriptional regulators. **a**, Morphological changes of EBs during hematopoietic differentiation. Representative NAMC microscopy images of EBs during differentiation are shown. The initial numbers of cells per EB for parameters #24, 107, 127, 167, 198, and 237 are 500, 800, 600, 600, 900, and 1,600, respectively. Flow cytometry plots of day 14 EBs are shown in Extended Data Fig. 6c. Scale bar, 200 µm. **b**, Immunofluorescence analysis of day 8 EBs cultured under the HPC and Hema conditions. Two representative maximum intensity projections of confocal microscopy images are shown for each condition. Inset panels show merged fluorescence and bright-field images. Scale bar, 100 µm.

These elongated EBs are reminiscent of gastruloids, hPSC-derived organoids that recapitulate some aspects of embryonic patterning during gastrulation^14^. To examine the formation of axial patterning, we performed whole-mount immunofluorescence staining on day 8 EBs. We observed that under the HPC conditions, a posterior endoderm and mesoderm marker CDX2 was localized at the smaller pole of elongated EBs, whereas endothelial cell markers CD34 and SOX17 were detected at the opposite pole, with CD34 localized in a vascular pattern (Fig. 6b and Extended Data Fig. 8a,b). RUNX1, a critical regulator of definitive hematopoiesis, was broadly expressed at the same pole but not in the same cells as SOX17. A few CD43^+^ cells were observed within a subset of RUNX1^+^ cells in a spatially random manner. GATA6 was expressed at the same pole but not in the same cells as CDX2, highlighting the difference from the previously reported gastruloid models with the anterior-posterior axis^14, 15^. EBs cultured under the Hema conditions did not exhibit the strong polarized expression of CDX2 and sometimes included more than one weak CDX2 expression domains. As a result, EBs were not elongated and the expression of SOX17 and RUNX1 was not polarized. Although an even larger number of CD43^+^ cells were observed at day 8 compared with the HPC conditions, these CD43^+^ cells did not necessarily appear to arise from RUNX1^+^ cells.

We also analyzed the day 8 EBs cultured in the absence of key signaling molecules. The EBs cultured without BMP4 also did not become elongated, yet CDX2 expression was detectable in at least one of the HPC conditions with larger EBs (Parameter ID #112; Extended Data Fig. 9). Removal of CHIR in M1 either in the presence or absence of Wnt-C59 in M2 resulted in the lack of robust CDX2 expression and the formation of irregularly shaped spherical aggregates. The EBs treated with an increased concentration of CHIR also failed to induce CDX2 expression and elongation under one of the HPC conditions (Parameter ID #24), indicating that an appropriate level of WNT/β-catenin signaling activity is required to robustly induce symmetry breaking and self-organized axial patterning. These findings indicate that robust and reproducible induction of HPCs from hPSCs leverages the design principles of cell fate decisions during embryonic development.

## Discussion

Developing robust and scalable differentiation protocols from hPSCs has been hampered by variability arising from biological and technical sources. These challenges often result in inconsistent cell populations and hinder reproducibility across laboratories and manufacturing sites, posing significant barriers to clinical translation. By integrating highly flexible robotic automation with AI-driven optimization, our study directly addresses common sources of variability through minimal modifications to existing manual experimental operations, enabling the rapid identification of culture conditions that robustly and reproducibly generate HPCs across multiple hiPSC lines. The highly efficient induction of HPCs from hiPSCs by our relatively simple three-step protocol would not only serve as a basis for more reliable and scalable manufacturing of immune cell therapeutics but also provide a valuable tool for understanding diseases associated with hematopoietic development or dysfunction.

Precise control over the dosage and timing of multiple signaling pathways enables robust and reproducible differentiation of hPSCs into HPCs through the formation of self-organized 3D structures with polarized expression of key transcriptional regulators required for endothelial and hematopoietic development. Stepwise regulation of signaling pathways—comprising an initial mesoderm induction phase with high FGF, intermediate WNT and BMP, and low Activin/Nodal signaling; a subsequent endothelial differentiation phase with high FGF and intermediate VEGF signaling accompanied by modest repression or downregulation of WNT and sufficient repression of Activin/Nodal signaling; and a final hematopoietic differentiation phase with downregulation of VEGF and activation of SCF signaling—is critical for efficient differentiation (Fig. 7). In particular, activation of WNT and FGF signaling in the initial phase favors the acquisition of HPC rather than Hema fate. Considering the presence of CD235a^+^ cells in Hema but not in HPC conditions, and in light of previous reports linking WNT and Activin/Nodal signaling to definitive and primitive hematopoiesis, respectively^6, 10^, HPCs and Hema might arise from the corresponding lineages. The EBs cultured under the HPC conditions activate CDX2 and show polarized expression of SOX17 and RUNX1 before generating CD43^+^ hematopoietic cells. This spontaneous symmetry breaking of an initially homogeneous cell population occurs in a highly reproducible manner within a specific 3D environment when exposed to a spatially uniform concentration of extrinsic diffusible factors. Symmetry breaking during development often relies on a combination of local activation via positive feedback and long-range inhibition mediated by rapidly diffusing inhibitors. Consistent with our observation that WNT and FGF signaling pathways play a key role, a recent study using a human gastruloid model of neural tube formation revealed that canonical WNT and FGF ligands, which act as slowly diffusing activators, and their feedback inhibitors, which function as rapidly acting diffusible or intracellular inhibitors that propagate their effects across neighboring cells, induce anterior-posterior symmetry breaking in a deterministic manner^16^. We also found that the exposure of EBs to intermediate levels of CHIR led to the robust induction of axial elongation and subsequent HPC generation. Such a moderate level of WNT activity appears to be optimal for initiating symmetry breaking, as it is high enough to activate WNT-dependent positive-feedback loops, while still maintaining a sufficient contrast between WNT-active and WNT-inactive cells in the presence of endogenous WNT ligands and inhibitors. This resulting heterogeneity in WNT activation may be sufficient to trigger self-organization through directed cell migration and sorting, ultimately resulting in the establishment of a single posterior pole of high WNT activity^17^. The requirement of exogenous ActA in this process was context-dependent.

**Fig. 7.**
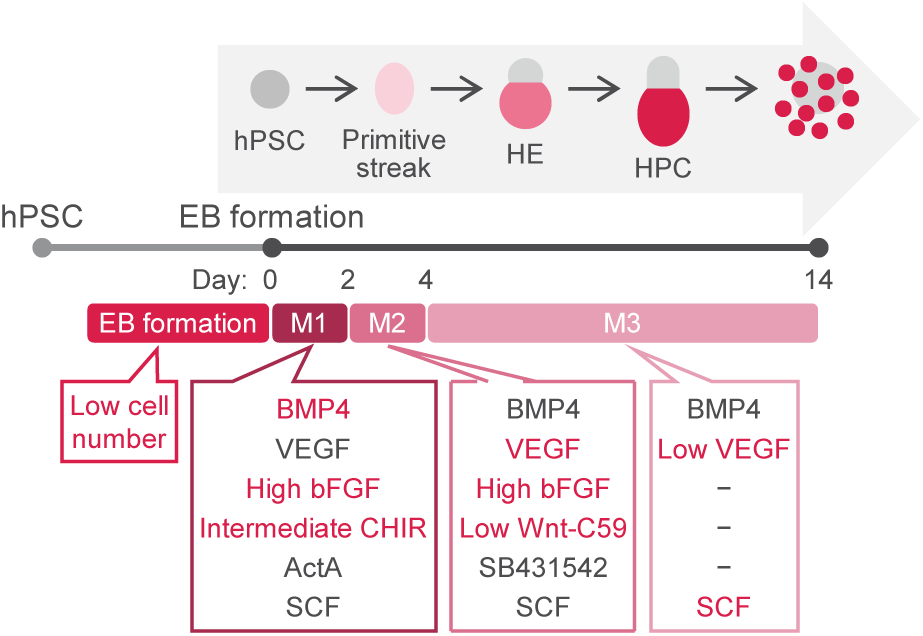
Key process parameters for the differentiation of HPCs from hPSCs. The critical process parameters identified and optimized are highlighted in red.

Although high Activin/Nodal signaling contributes to the acquisition of Hema rather than HPC fate, the addition of exogenous ActA in one of the HPC conditions with smaller EBs was necessary for consistent EB growth and development. Interestingly, a study using a mouse gastruloid model revealed that endogenous Nodal activity increased in proportion to initial aggregate size^17^. As increased EB size is associated with inefficient differentiation into HPCs in our system, supplementation of ActA in smaller aggregates may simply compensate for their lower endogenous Nodal levels, thereby supporting adequate growth without driving them toward non-HPC fates.

Our unbiased and systematic robotic search enabled a marked improvement in differentiation efficiency by simultaneously optimizing multiple signaling inputs. Interpreting these optimized conditions within a developmental framework suggests that such automated approaches can also provide useful clues to the underlying biology. We anticipate that combining advanced automation technologies with human insight will transform research and development of stem cell therapies.

## Methods

### Maholo robotic platform

Our robotic platform “Maholo” (Robotic Biology Institute) was designed to automate a variety of cell culture processes using a programmable robot. It consists of a dual-arm robot (MOTOMAN-CSDA10F, YASKAWA Electric) that can imitate natural human motions and operate a number of common laboratory instruments and pieces of equipment such as incubators (SCALE, RORZE Lifescience), centrifuges, microscopes (BioStudio-T with phase-contrast and BioStudio-T with Nikon Advanced Modulation Contrast [NAMC], Nikon), cell counters (Cellaca MX, Revvity), refrigerators, dry baths, micropipettes and aspirators.

### hiPSC culture

An in-house hiPSC line^18^ (dermal fibroblast-derived) and three hiPSC lines obtained from RIKEN BRC (peripheral blood-derived 1383D6 and 1231A3, and dermal fibroblast-derived 409B2) were maintained on iMatrix-511 (Matrixome)-coated plates in StemFit AK02N medium (Ajinomoto) at 37 °C in 5% CO2 in air.

### EB formation using the robotic automation

For EB formation, hiPSCs were manually dissociated into single cells using Accutase (Innovative Cell Technologies), stained with Acridine Orange and DAPI, and viable cells were counted using an automated cell counter (NucleoCounter NC-250, ChemoMetec). After manual adjustment of cell concentration, Maholo seeded defined numbers of viable cells into AggreWell 800 24-well plates (STEMCELL Technologies) containing approximately 300 microwells per well. The plates were left undisturbed at ambient temperature for at least 60 min before being returned to the 37 °C incubator with 5% CO2 in air. Medium was changed by Maholo every two to three days by removing half of the spent medium and replacing it with an equal volume of fresh medium.

To induce stepwise differentiation of hiPSCs into posterior primitive streak/lateral plate mesoderm, endothelial cells and HPCs through temporally controlled exposure to signaling cues, EBs were cultured sequentially in three different media over a 14-day period as follows:

Mesoderm induction medium (M1; from day 0 to day 2) — EB formation medium supplemented with 10 µM Y-27632, 20 ng/mL SCF (PeproTech), and various concentrations of BMP4 (R&D Systems), bFGF (PeproTech), CHIR99021 (CHIR), Activin A (ActA; R&D Systems), and VEGF (PeproTech).

Endothelial differentiation medium (M2; from day 2 to day 4) — EB formation medium supplemented with 10 ng/mL BMP4, 10 ng/mL SCF, and various concentrations of bFGF, Wnt-C59, SB431542, and VEGF.

Hematopoietic differentiation medium (M3; from day 4 to day 14) — EB formation medium supplemented with 10 ng/mL BMP4 and various concentrations of VEGF and SCF.

The EB formation medium consists of STEMdiff APEL2 Medium (APEL2; STEMCELL Technologies) supplemented with 5% Protein-Free Hybridoma Medium II (PFHM II; Gibco). Parameter values are given as final concentrations in 2 mL of culture medium. The amount of cytokines and chemical compounds remaining in the culture medium at the time of medium change, which should be gradually degraded or diluted by subsequent medium changes, was not taken into account in the calculations. Half-medium changes were performed on day 2 with the endothelial differentiation medium and on days 4, 7, 10, and 12 with the hematopoietic differentiation medium using Maholo equipped with automatic micropipettes and taking care not to disturb the settled EBs. Specifically, half (1 mL) of the culture medium was removed from each well at an aspiration rate of 232 µL/sec, various volumes (5–100 µL) of each stock solution were added at a dispense rate of 36 µL/sec, and EB formation medium was added to adjust the total volume to 2 mL at a dispense rate of 47 µL/sec. Medium changes were carried out by removing one plate at a time from the incubator, with a process time limited to 30-45 min per plate.

In the standard condition, hiPSCs were seeded at 3,000 cells per microwell (9×10^5^ cells per well of an AggreWell 800 24-well plate) and treated with 20 ng/mL BMP4, 20 ng/mL VEGF, and 20 ng/mL SCF for two days, followed by 10 ng/mL BMP4, 20 ng/mL VEGF, and 10 ng/mL SCF for two days, then 10 ng/mL BMP4, 20 ng/mL VEGF, and 20 ng/mL SCF for another 10 days. The medium supplied on day 0 also contained 10 µM Y-27632.

Process parameters for EB formation and HPC differentiation were optimized using Epistra Accelerate v2.0.0 (Epistra Inc.; https://www.epistra.jp/epistra-accelerate/docs/), a batch BO software designed for life science applications. The software constructs a Gaussian process surrogate model that jointly predicts experimental outcomes and quantifies predictive uncertainty across the parameter space. By leveraging this uncertainty estimate, the algorithm proposes batches of candidate conditions that balance exploitation of regions predicted to yield favorable outcomes with exploration of regions where the model remains uncertain, thereby efficiently navigating high-dimensional search spaces within a limited number of experimental iterations. The algorithms implemented in the software have been designed and tested on benchmark problems that recapitulate real-world cell culture media optimization tasks.

Phase-contrast or NAMC microscopy images were acquired using BioStudio-T microscopes integrated into the Maholo platform. Multiple images were stitched together to generate large field-of-view images using the software’s built-in tool.

### Flow cytometry

For flow cytometry of dissociated EBs, all cells derived from EBs were manually harvested from microwell plates on day 14 and digested with 2,500 U/mL Collagenase II (STEMCELL Technologies) at 37 °C for 20 min, then treated with Accumax (Innovative Cell Technologies) at 37 °C for 10 min, followed by repeated pipetting and an additional 10-min Accumax treatment to maximize single-cell dissociation. Viable cell numbers were determined for a subset of samples. Cells were stained with Zombie Violet fixable viability dye (BioLegend) and subsequently incubated with human Fc block (BD Biosciences), Super Bright Complete Staining buffer (eBioscience), and antibodies against CD90/THY1, CD34, CD43/SPN, CD45/PTPRC, CD235a/GYPA, and lineage markers (CD3, CD14, CD16, CD19, CD20, and CD56). Samples were filtered through a 70 µm mesh and analyzed using a NovoCyte Advanteon flow cytometer (Agilent) equipped with three lasers (405 nm, 488 nm, and 640 nm). Fluorescence compensation was performed for each experiment using single-stained compensation beads (ArC amine reactive beads [Thermo Fisher] or UltraComp eBeads [Thermo Fisher]). Debris, doublets, and dead cells were excluded based on forward/side scatter and Zombie Violet staining (Extended Data Fig. 2a). Lineage-positive cells were excluded using the same channel (V445) as that used for Zombie Violet. Unstained and/or isotype controls were used to assess non-specific staining.

For flow cytometry of NK cells, cells in suspension were harvested and the number of viable cells was counted. Cells were stained with LIVE/DEAD Fixable Near-IR Stain (Thermo Fisher), followed by incubation with human Fc block and antibodies against CD56/NCAM1, CD45/PTPRC, CD226/DNAM-1, CD16/FcγRIII, CD314/NKG2D, CD94/NKG2A, CD336/NKp44, CD335/NKp46, and CD3.

The list of antibodies used for flow cytometry is shown in Supplementary Table 4.

### Computational analysis of flow cytometry data

Flow cytometry data were preprocessed in R using the flowCore (v2.18.0) and flowWorkspace (v4.18.0) packages. Debris, doublets, dead cells, and lineage-positive cells were manually excluded through sequential gating. The compensated fluorescence intensity values were transformed using the logicle transformation with identical parameter settings across all experiments, and then Z-score normalized for each marker. For hematopoietic score calculation, Z-score normalization was performed at the start of each optimization iteration using all data generated up to that iteration. For subpopulation analysis, normalization parameters were computed from the full dataset.

For dimensionality reduction, up to 1,000 gated cells were randomly selected from each sample, and UMAP analysis was performed based on the expression levels (peak height) of CD90, CD34, CD43, CD45, and CD235a using the umap package (v0.2.10.0) with n_neighbors = 10 and min_dist = 0.1. Clustering was then performed using the dbscan package (v1.2-0) implementing HDBSCAN with minPts = 13. A broad cluster with CD43 expression was annotated as hematopoietic cells. The percentage of viable cells belonging to the CD43^+^ cluster (“a hematopoietic score”) was quantified for each sample and used as the objective function for optimization, given that CD34^+^/CD43^+^/CD45^+^ HPCs are thought to exist only transiently during hPSC differentiation before further differentiating into CD34^−^ hematopoietic cells. UMAP was recomputed after the incorporation of newly obtained data following the second iteration, whereas data obtained in the third iteration were projected onto the previously computed UMAP embedding.

To model the relationship between multiple process parameters and differentiation outcomes, we performed Partial Least Squares (PLS) analysis, a multivariate regression approach that identifies latent variables representing weighted combinations of process parameters maximizing covariance with the hematopoietic score.

To identify different subpopulations of hematopoietic cells, FlowSOM (v2.14.0) analysis was performed based on the signal pulse areas using the same set of 1,000 live single-cell events per sample that were used for hematopoietic score calculation. These events were assigned to a self-organizing map (SOM) with a 7 × 7 grid (xdim = 7, ydim = 7), resulting in a minimum spanning tree with 49 nodes. Similar SOM nodes were merged into 15 biologically meaningful metaclusters using the consensus clustering algorithm implemented in FlowSOM.

To cluster the samples based on their cell population distributions, the proportion of cells in each metacluster was calculated for each sample, generating compositional profiles. Pairwise similarities in cell-type distributions between samples were quantified based on the Jensen–Shannon distance (JSD) using the philentropy package (v0.9.0), and the resulting distance matrices were used for clustering with the partitioning around medoids (PAM) algorithm from the cluster package in R. These analyses were performed on all 300 samples, as well as on a subset of 137 samples with the hematopoietic score ≥ 0.75.

A random forest classifier was trained to distinguish the HPC group from other phenotypic groups based on process parameter values from the 137 samples with the hematopoietic score ≥ 0.75. Model hyperparameters were optimized by 10-fold cross-validation to balance classification accuracy and receiver operating characteristic (ROC) area under the curve (AUC), yielding an accuracy of 0.906 and a ROC AUC of 0.971. The final model was trained on the full set of 137 samples using the tuned hyperparameters. SHapley Additive exPlanations (SHAP) values were computed for each sample to assess the magnitude and direction of individual parameter contributions to the model’s predictions.

### NK cell differentiation

For NK cell differentiation, all cells derived from EBs were manually harvested from the microwell plates on day 14 and triturated by pipetting. Subsequently, the cells were collected by centrifugation, resuspended in NK cell differentiation medium, and seeded into non-TC-treated 6-well plates precoated with 0.5 µg/cm^2^ human DLL4-Fc (ACRO Biosystems) and 0.5 µg/cm^2^ human fibronectin fragment RetroNectin (Takara Bio) at 37 °C for 2 h. One-tenth of the cells from each well of AggreWell 800 24-well plates were seeded into a well of a six-well plate for NK cell differentiation and the remaining cells were analyzed by flow cytometry. NK cell differentiation medium consists of a 2:1 mixture of DMEM (Gibco) and Ham’s F-12 (Gibco) supplemented with 7.5% heat-inactivated human AB serum (BioIVT), 5.0 ng/mL sodium selenite (FUJIFILM Wako), 50 µM ethanolamine (Merck Millipore), 32.9 µg/mL L-ascorbic acid 2-phosphate (Sigma-Aldrich), 20 ng/mL SCF (PeproTech), 20 ng/mL IL-7 (PeproTech), 10 ng/mL IL-15 (PeproTech), 10 ng/mL FLT3L (PeproTech), and 35 nM UM171. Half of the medium was replaced with fresh medium two to three times a week. After 2–3 weeks of NK cell differentiation, the cells were harvested and either seeded for NK cell maturation or cryopreserved in STEM-CELLBANKER (Takara Bio) or CryoStor CS10 (STEMCELL Technologies) freezing media.

For NK cell maturation, the cells were cultured in CTS NK-Xpander Medium (Thermo Fisher) supplemented with 7.5% heat-inactivated human AB serum, 10 ng/mL IL-15, 10 ng/mL IL-18 (R&D Systems), and 10 ng/mL IL-21 (PeproTech) for three days on non-TC-treated plates precoated with 0.76 µg/cm^2^ anti-NKp30 antibody (BioLegend), 0.76 µg/cm^2^ anti-DNAM-1 antibody (BioRad), and 0.76 µg/cm^2^ human fibronectin fragment RetroNectin overnight at 4 °C. The cells were then transferred to stirred bioreactors (BWV-S03A, ABLE-Biott) and cultured in CTS NK-Xpander Medium supplemented with 7.5% heat-inactivated human AB serum, 10 ng/mL IL-15, and 10 ng/mL IL-18 at 55 rpm in a 37 °C incubator with 5% CO2 for three days. The cells were harvested and either used for cytotoxicity assays or analyzed by flow cytometry.

### Cytotoxicity assay

The natural cytotoxicity of NK cells was measured by co-culturing NK cells with K562-HiBiT cells (Promega) at various effector-to-target (E:T) ratios in 96-well U-bottom plates in triplicate. After four hours of incubation, HiBiT-tagged proteins released from lysed cells were detected using Nano-Glo HiBiT Extracellular Detection System (Promega). The luminescent signal was measured on a plate reader. The percentage of specific lysis was calculated as follows: %Specific lysis = (Bioluminescence in the test samples − Background signal in the absence of effector cells)/(Bioluminescence in the presence of digitonin − Background signal in the absence of effector cells) × 100.

### Cytokine secretion assay

To assess the capacity of IFN-γ and TNF-α secretion upon stimulation with tumor cells, NK cells were co-cultured with K562 cells at 1:1 E:T ratio in 96-well U-bottom plates in triplicate. After 24 hours of incubation, culture supernatants were harvested for IFN-γ and TNF-α measurements with AlphaLISA Cytokine Kits (Revvity).

### CFU assay

For the CFU assay, the EB-derived, singly dissociated cells were resuspended in methylcellulose media (MethoCult H4034 Optimum, STEMCELL Technologies) and seeded in duplicate at 2,000 cells/well into non-TC-treated 6-well plates. On day 15, the entire wells were imaged using a Nikon Eclipse Ti2 microscope and colonies were scored for colony-forming unit erythroid (CFU-E), burst-forming unit erythroid (BFU-E), CFU-G (granulocyte), CFU-M (macrophage), CFU-GM (granulocyte-macrophage), and CFU-GEMM (granulocyte, erythrocyte, macrophage, megakaryocyte) on stitched images.

### Immunofluorescence

EBs were harvested from the microwell plates on day 8 and fixed in 4% paraformaldehyde at 4 °C for 2 h. After permeabilization and blocking with 0.2% Triton X-100 and 10% fetal bovine serum, EBs were incubated with primary antibodies overnight at 4 °C. After washing, EBs were incubated with fluorochrome-conjugated secondary antibodies and 1 µg/mL Hoechst 33342 overnight at 4 °C. After washing, EBs were mounted on microscope slides with SlowFade Glass Soft-set Antifade Mountant (Thermo Fisher). Images were acquired using a CQ1 spinning disk confocal system (Yokogawa Electric Corporation). The list of primary and secondary antibodies used for immunofluorescence is shown in Supplementary Table 5.

## Acknowledgments

This work was funded by Astellas Pharma Inc. We thank Yosuke Ozawa, Taku Tsuzuki, Tsuyoshi Tatsukawa, Tatsuki Higashi, and Kensuke Tozuka (Epistra Inc.) for providing the proprietary BBO platform *Epistra Accelerate* and for their thoughtful advice and critical review of the manuscript. We are also grateful to Robotic Biology Institute Inc. for technical assistance with the robotic system and RIKEN BRC for providing the hiPSC lines. We appreciate Tohru Natsume (The National Institute of Advanced Industrial Science and Technology) for helpful discussions, Hisayo Yamaguchi and Yuko Oda for technical assistance, and Masazumi Kamohara and Shinsuke Oshima for their critical review of the manuscript.

## Author contributions

K.A., A.I., and H.Y. conceived the study. K.A. designed the study. K.A., N.O., Y.G., F.S., and M.I. performed the experiments. K.A., A.S., and M.I. analyzed the data. Y.T. provided technical support. K.A. wrote the manuscript, and A.S. and M.I. reviewed and edited the manuscript. All authors reviewed and approved the final manuscript.

## Competing interests

K.A., N.O., Y.G., Y.T., A.I., and H.Y. are employees of Astellas Pharma Inc. and Cellafa Bioscience Inc. A.S., F.S., and M.I., are employees of Astellas Pharma Inc.

## Supplementary Information

**Extended Data Fig. 1.**
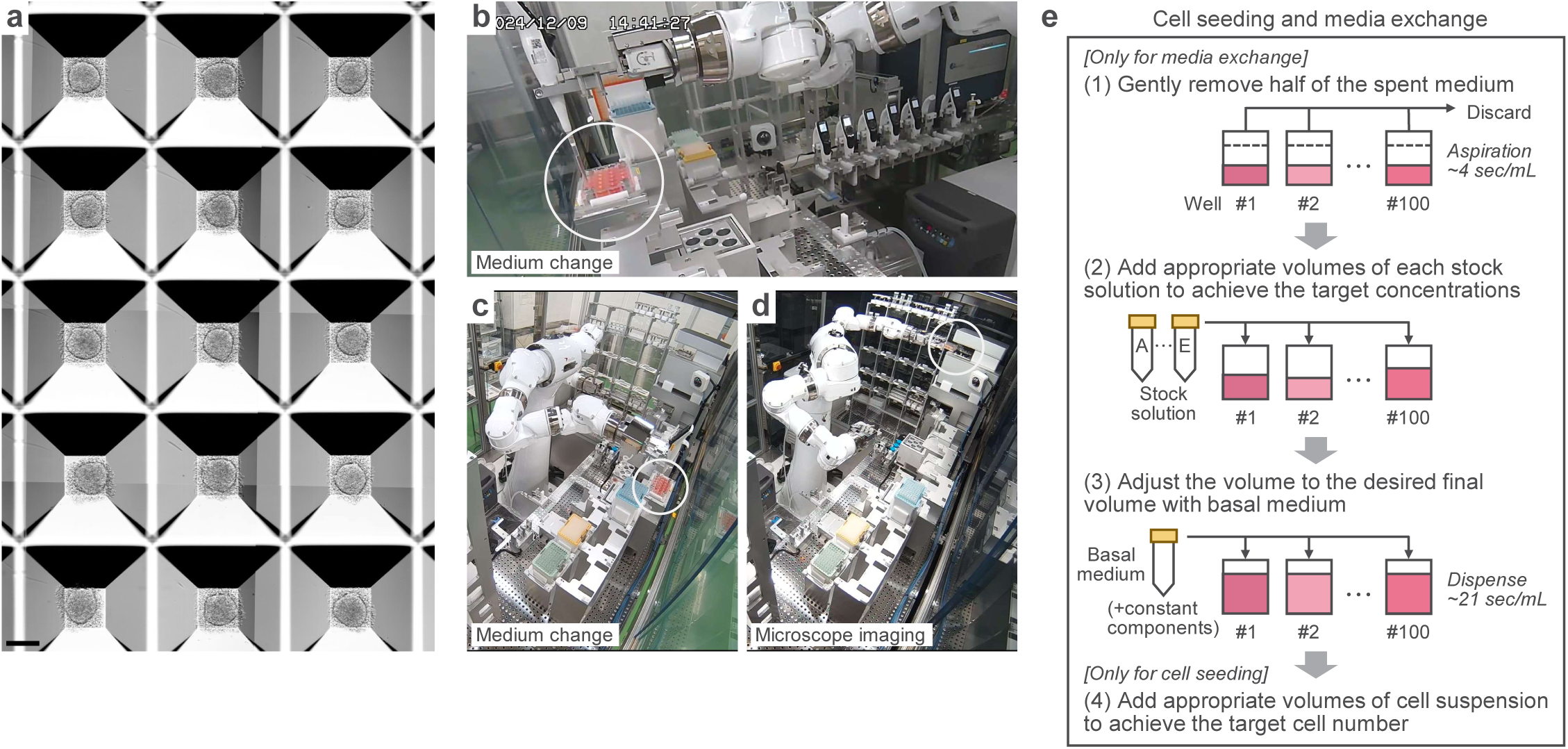
The automated workflow enabling parallel testing of multiple conditions for EB formation. **a**, EBs formed in microwells one day after seeding. Multiple images were acquired with a Nikon Advanced Modulation Contrast (NAMC) microscope integrated into the robotic platform and were stitched together to produce a large field-of-view image. Scale bar, 200 µm. **b-d**, Robot performing medium changes (**b** and **c**) and microscope imaging (**d**). **e**, A schematic illustration of the procedure of cell seeding and media exchanges. To generate EBs under various conditions with different numbers of cells per EB and varying concentrations of signaling molecules, stock solutions of cells, cytokines, and chemical compounds were manually prepared, and Maholo was programmed to dispense specified volumes of each stock solution into the corresponding wells of microwell plates, thereby achieving the desired final concentrations. The addition of cell suspension (step 4) was performed exclusively for cell seeding, whereas medium removal (step 1) was performed exclusively during media exchanges. Cell seeding and media exchanges were carried out at ambient temperature outside the incubator, requiring approximately 30–45 min per plate.

**Extended Data Fig. 2.**
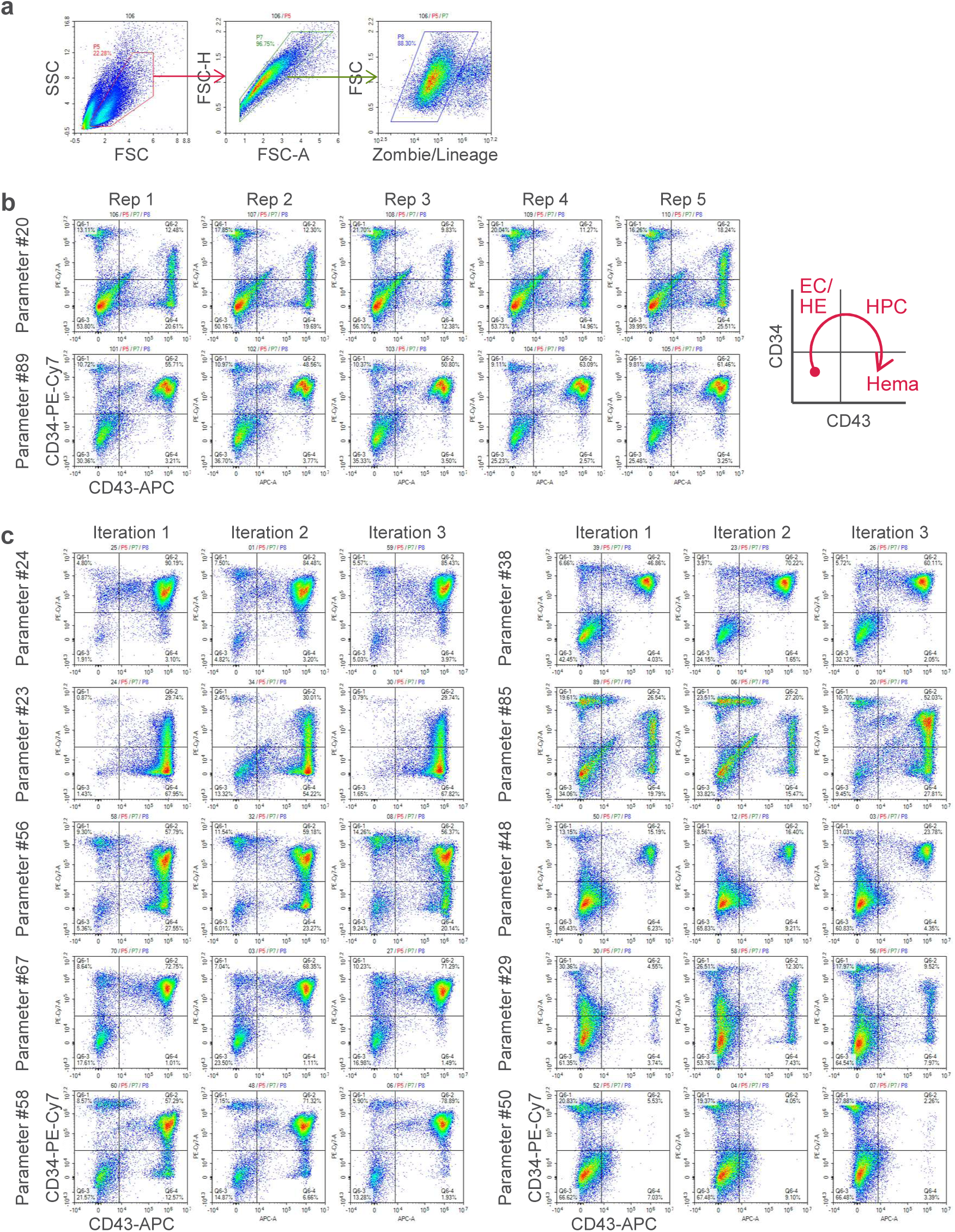
Reproducibility of differentiation assessed by flow cytometry. **a**, Representative gating strategy for flow cytometry analysis. Debris, doublets, dead cells, and lineage-positive cells were excluded by gating. **b**, Flow cytometry plots showing CD43 and CD34 expression from five replicates for each of two conditions within a single experiment. **c**, Representative flow cytometry plots for CD43 and CD34 expression in cell populations generated under identical conditions across three independent experiments. Only the first replicate is shown for each condition and experiment.

**Extended Data Fig. 3.**
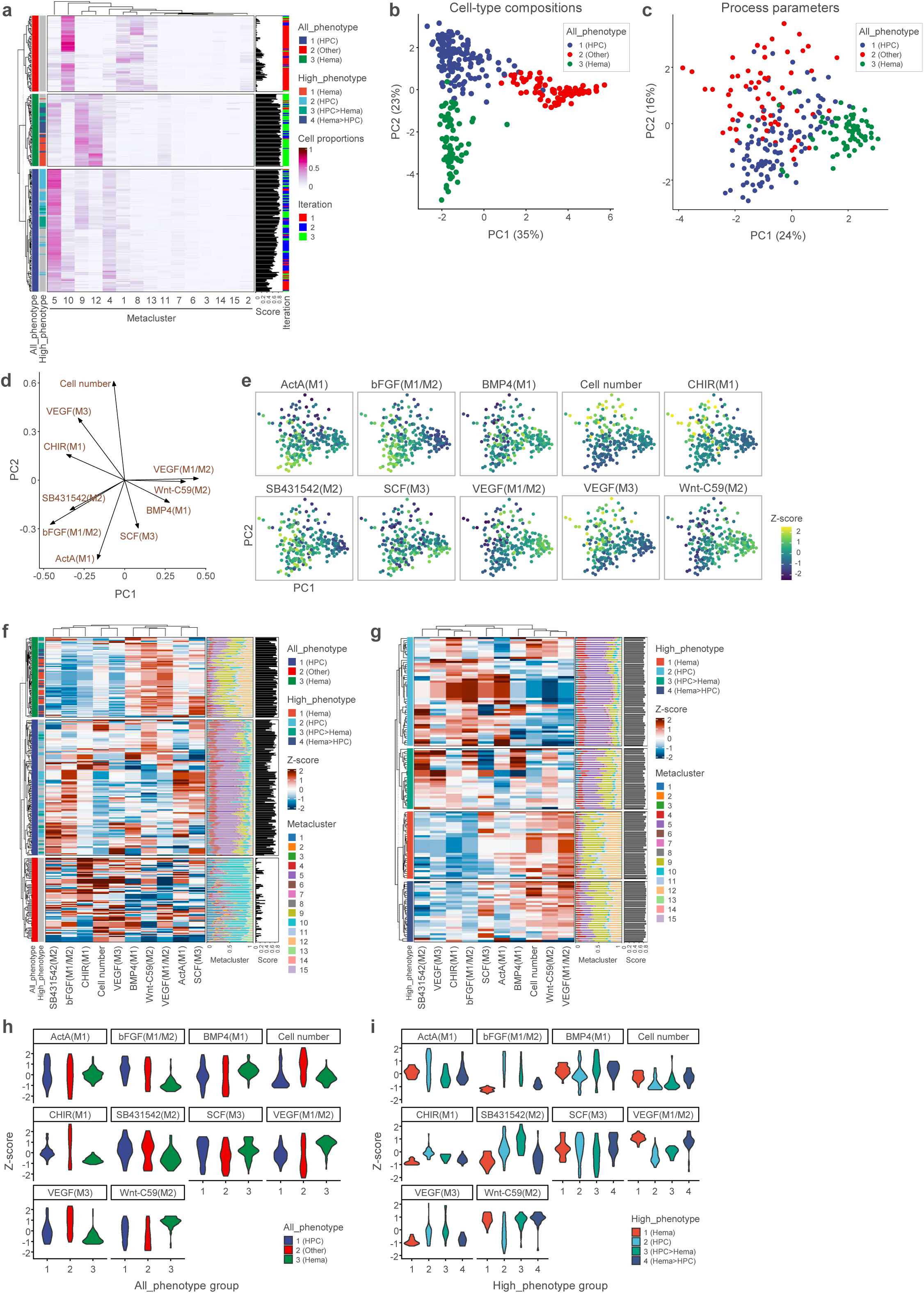
Multivariate characterization of differentiation outcomes using experimental conditions and cellular phenotypes. **a**, Heatmap showing compositions of metaclusters for all 300 samples. Three phenotypic groups were identified by PAM clustering based on JSD between samples. **b**, PCA plot based on compositions of metaclusters for all 300 samples. **c**, PCA plot based on process parameters. **d**, Loading plot of 10 process parameters for the first and second principal components. The direction and length of the vectors represent the contribution of each variable to the two principal components. **e**, PCA plot of process parameters with colors representing Z-score-normalized parameter values. **f**, Heatmap of process parameters for all 300 samples. Replicate samples under the same conditions are treated as independent samples. The proportions of cells in each metacluster and hematopoietic scores are shown next to the heatmap. **g**, Heatmap of process parameters for 137 high-score samples. Distinct patterns of elevated and reduced parameter values are observed across specific phenotypic groups. **h**, Violin plots of process parameters for each phenotype group. **i**, Violin plots of process parameters for each high-score phenotype group.

**Extended Data Fig. 4.**
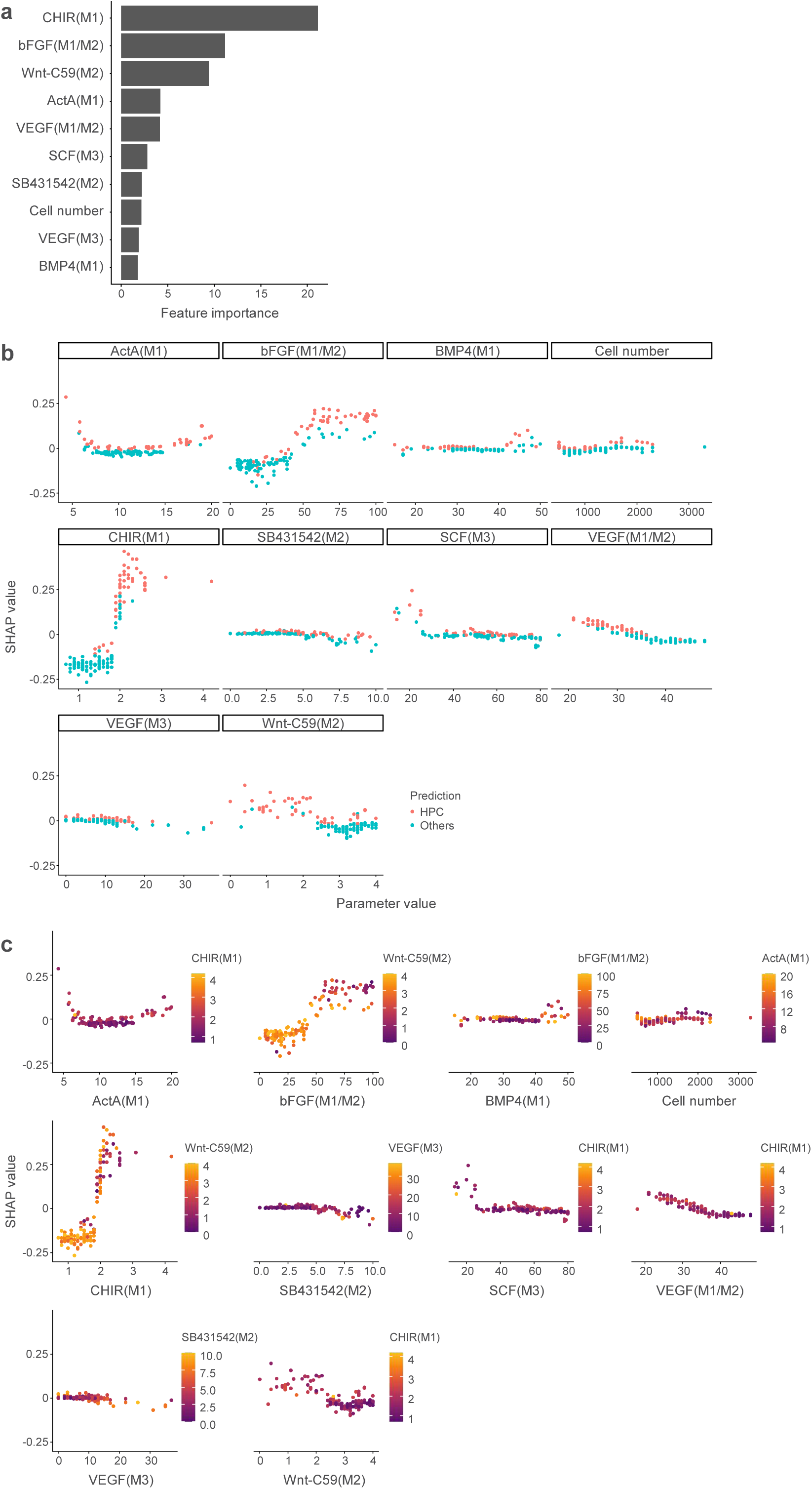
Feature importance in random forest classifier. **a**, Feature importance of the random forest model to predict differentiation outcomes. **b**, SHAP dependence scatter plot showing the impact of a single feature on the predictions of HPC fate. Lower CHIR in M1 and lower bFGF in M1/M2 are associated with a sharp decrease in SHAP values for the prediction of HPC fate. **c**, SHAP scatter plot highlighting interaction effects between features. The vertical dispersion in SHAP values indicates that the same value for a given feature can have a different impact on the model’s output depending on other features. A distinct vertical pattern of coloring suggests the interaction effects between bFGF in M1/M2 and Wnt-C59 in M2. Under the conditions with higher bFGF in M1/M2, a higher level of Wnt-C59 in M2 is associated with the decrease in SHAP values for this region.

**Extended Data Fig. 5.**
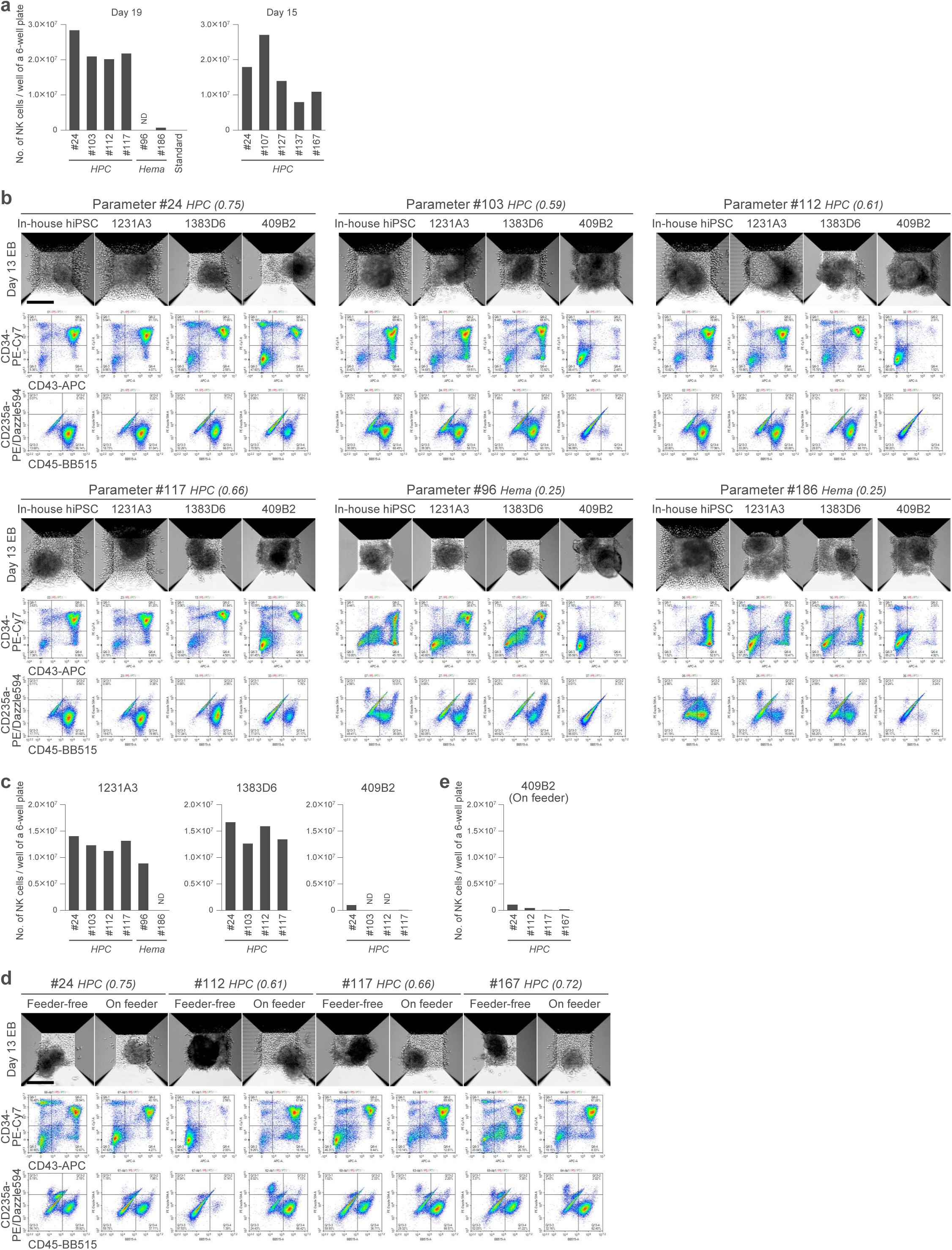
Generation of HPCs and NK cells from multiple iPSC lines. **a**, The number of NK cells obtained on day 19 (left) and day 15 (right) of differentiation from in-house iPSCs in a single well of a six-well plate, based on results from two additional experiments. ND, not determined due to poor growth. **b**, Representative NAMC microscopy images (top) and flow cytometry plots (middle and bottom) of day 14 EBs generated from multiple iPSC lines. Phenotype groups (1: HPC, 2: other, 3: Hema) and proportions of HPC are shown for each parameter ID. Scale bar, 200 µm. **c**, The number of NK cells obtained on day 19 of differentiation from 1383D6, 1231A3, and 409B2 iPSCs in a single well of a six-well plate. The data for in-house iPSCs are shown in **a** (left). ND, not determined due to poor growth. **d–e**, 409B2 iPSCs either cultured in feeder-free conditions or transferred onto human dermal fibroblast (HDF) feeder cells prior to EB formation were differentiated under HPC conditions. A short-term culture on feeders increased the differentiation potential of 409B2 iPSCs into CD34^+^/CD43^+^ cells in some conditions (**d**), yet these cells were not readily generated NK cells (**e**). The data for in-house iPSCs are shown in **a** (right). Scale bar in **d**, 200 µm.

**Extended Data Fig. 6.**
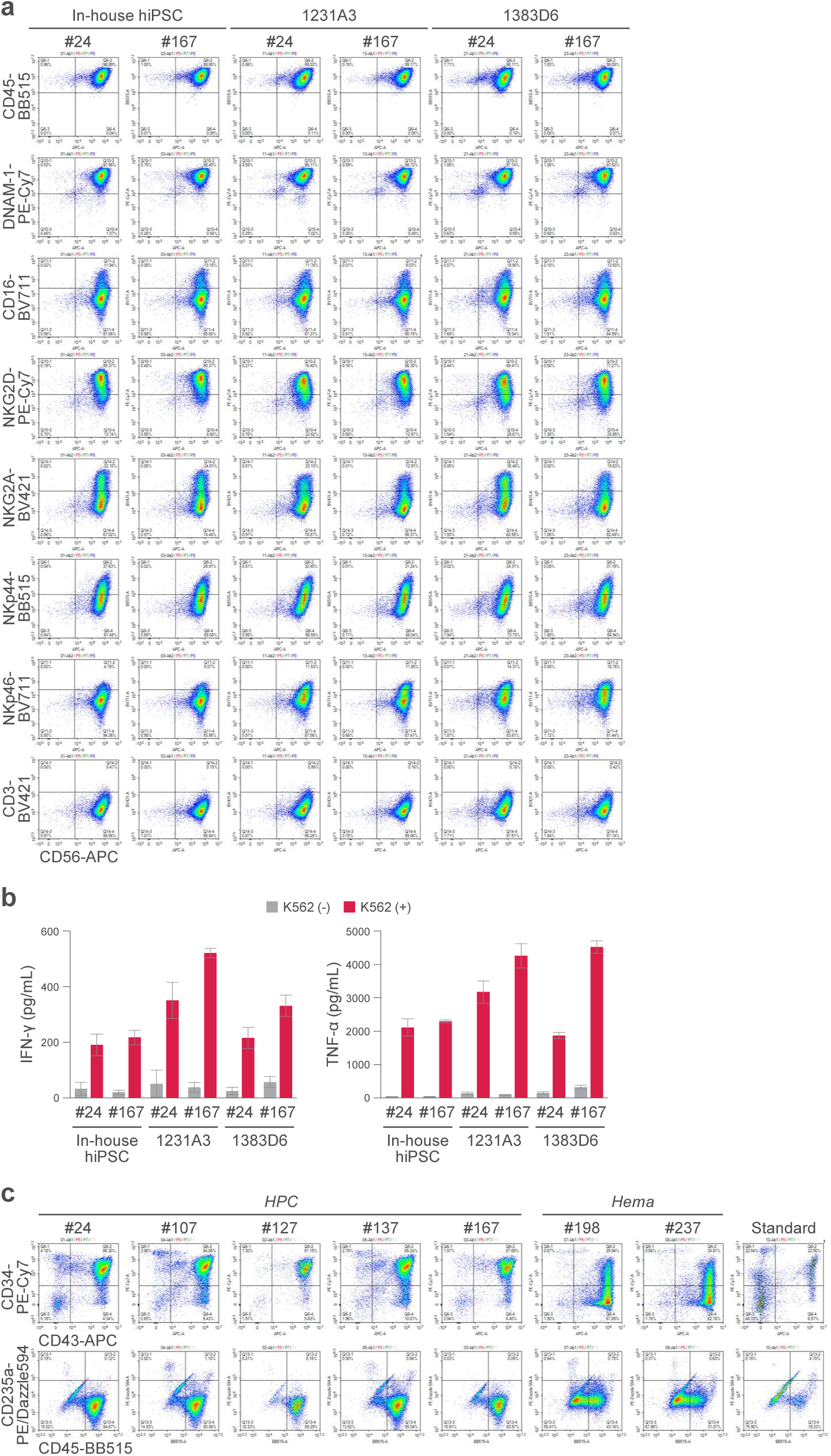
Phenotypic and functional characterization of hiPSC-derived NK cells. **a**, Flow cytometry analysis of mature NK cells. Three hiPSC lines were differentiated into HPCs under two different conditions for 14 days. HPCs were further cultured under NK cell differentiation conditions for 15 days, followed by NK cell maturation conditions for 6 days. **b**, IFN-γ and TNF-α secretion by NK cells after stimulation by K562 cells. Data represent mean ± SD of triplicates. **c**, Flow cytometry plots of day 14 EBs, from which samples for the CFU assays were derived. A smaller number of cells were analyzed for #127, 167, and the standard condition.

**Extended Data Fig. 7.**
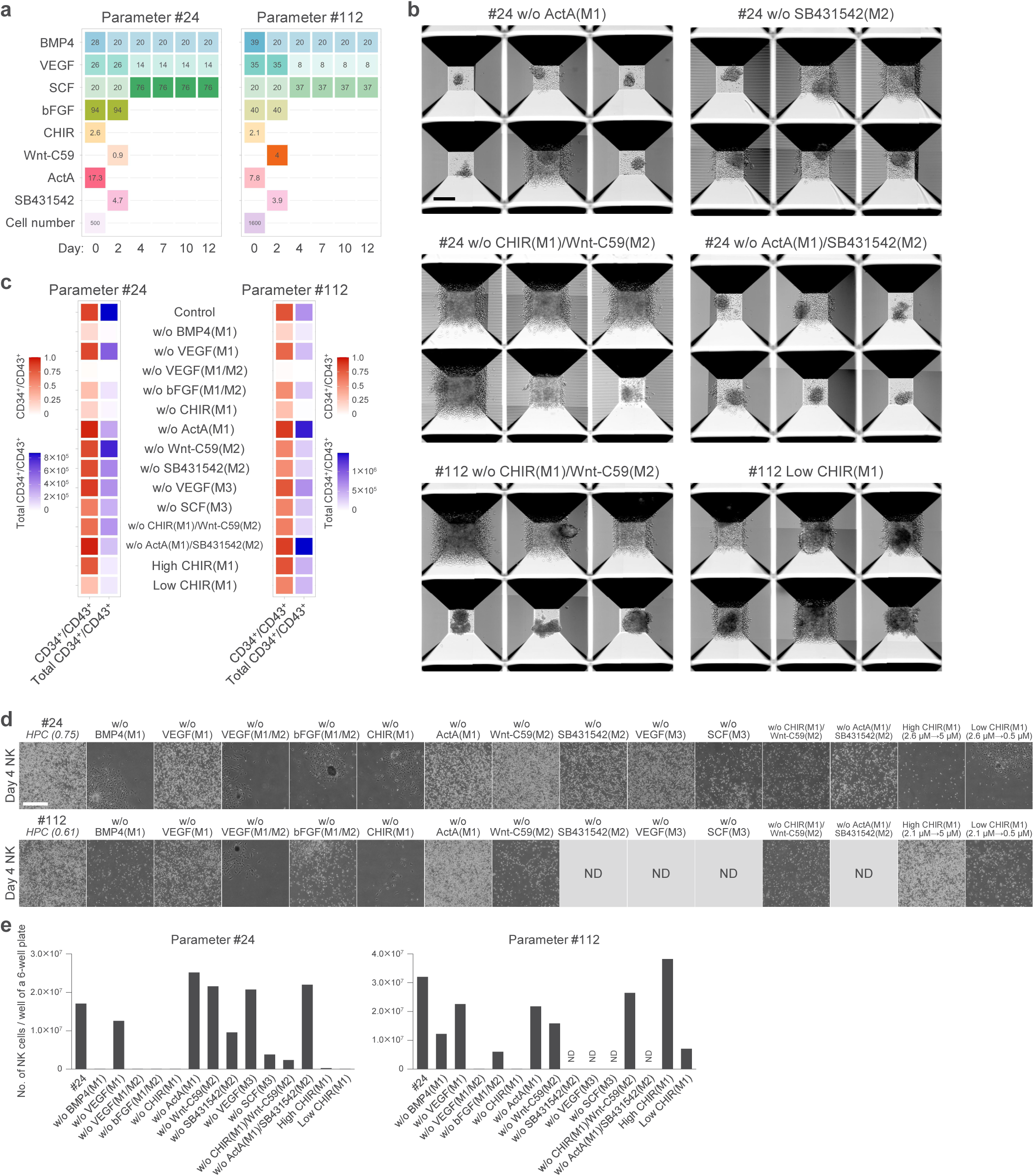
Dependence of HPC differentiation on a combination of key signaling pathways. **a**, Parameter values for two different HPC conditions. Values are expressed as the final concentrations of cytokines and chemical compounds in 2 mL of culture medium. **b**, Representative NAMC microscopy images of day 14 EBs from samples with high variability in EB morphology. Photographs were cropped from larger stitched images. Scale bar, 200 µm. **c**, Heatmaps showing the proportion and total number of CD34^+^/CD43^+^ cells per well of an AggreWell 800 24-well plate on day 14. **d**, Representative images of cells four days after seeding under NK cell differentiation conditions. ND, not determined. Scale bar, 500 µm. **e**, The number of NK cells obtained on day 15 of differentiation from a single well of a six-well plate. ND, not determined.

**Extended Data Fig. 8.**
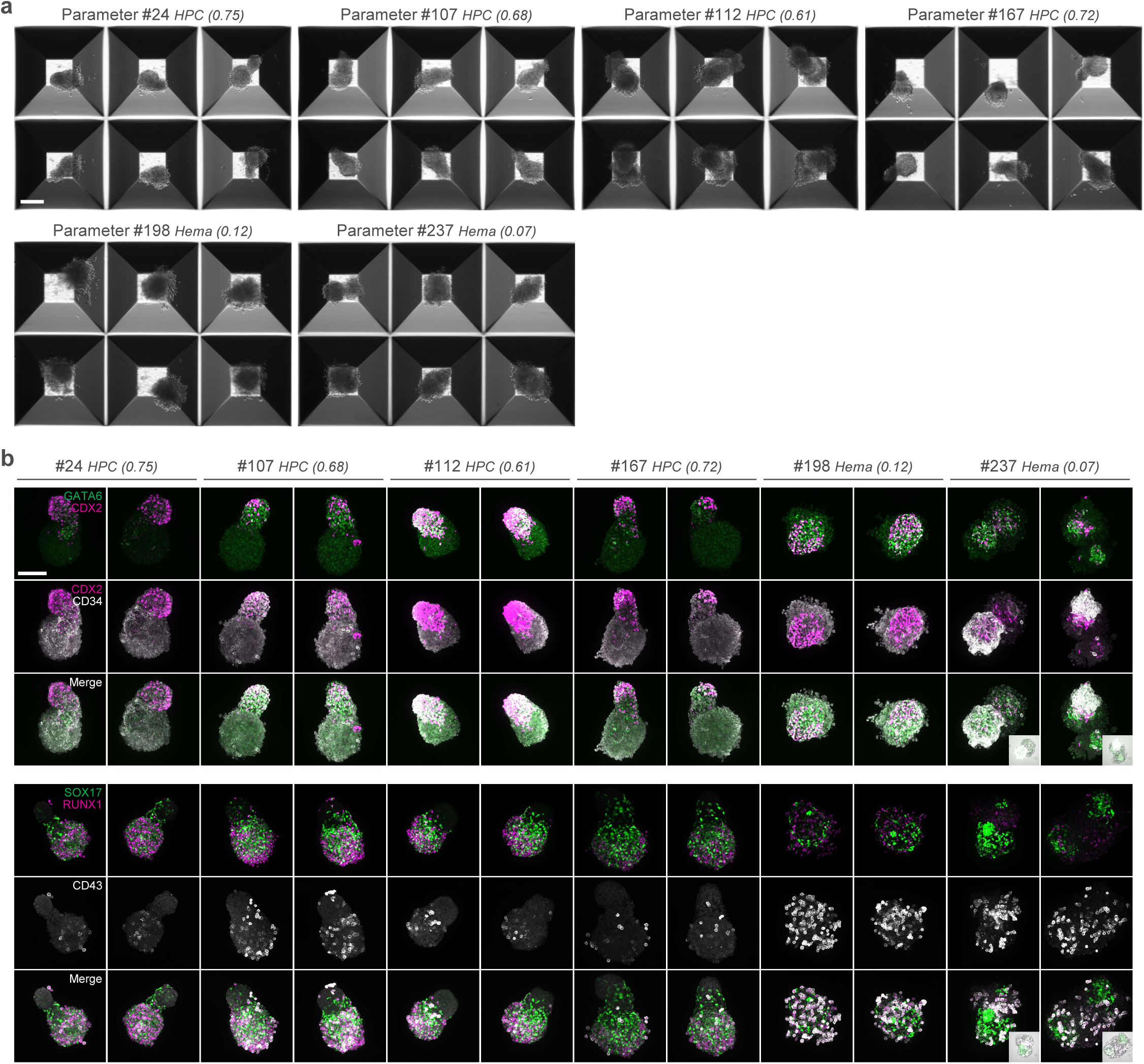
Spontaneous symmetry breaking in EBs prior to the emergence of HPCs. **a**, Representative NAMC microscopy images of day 8 EBs. Scale bar, 200 µm. **b**, Immunofluorescence analysis of additional marker combinations in day 8 EBs cultured under the HPC and Hema conditions. Two representative maximum intensity projections of confocal microscopy images are shown for each condition. Inset panels show merged fluorescence and bright-field images. Scale bar, 100 µm.

**Extended Data Fig. 9.**
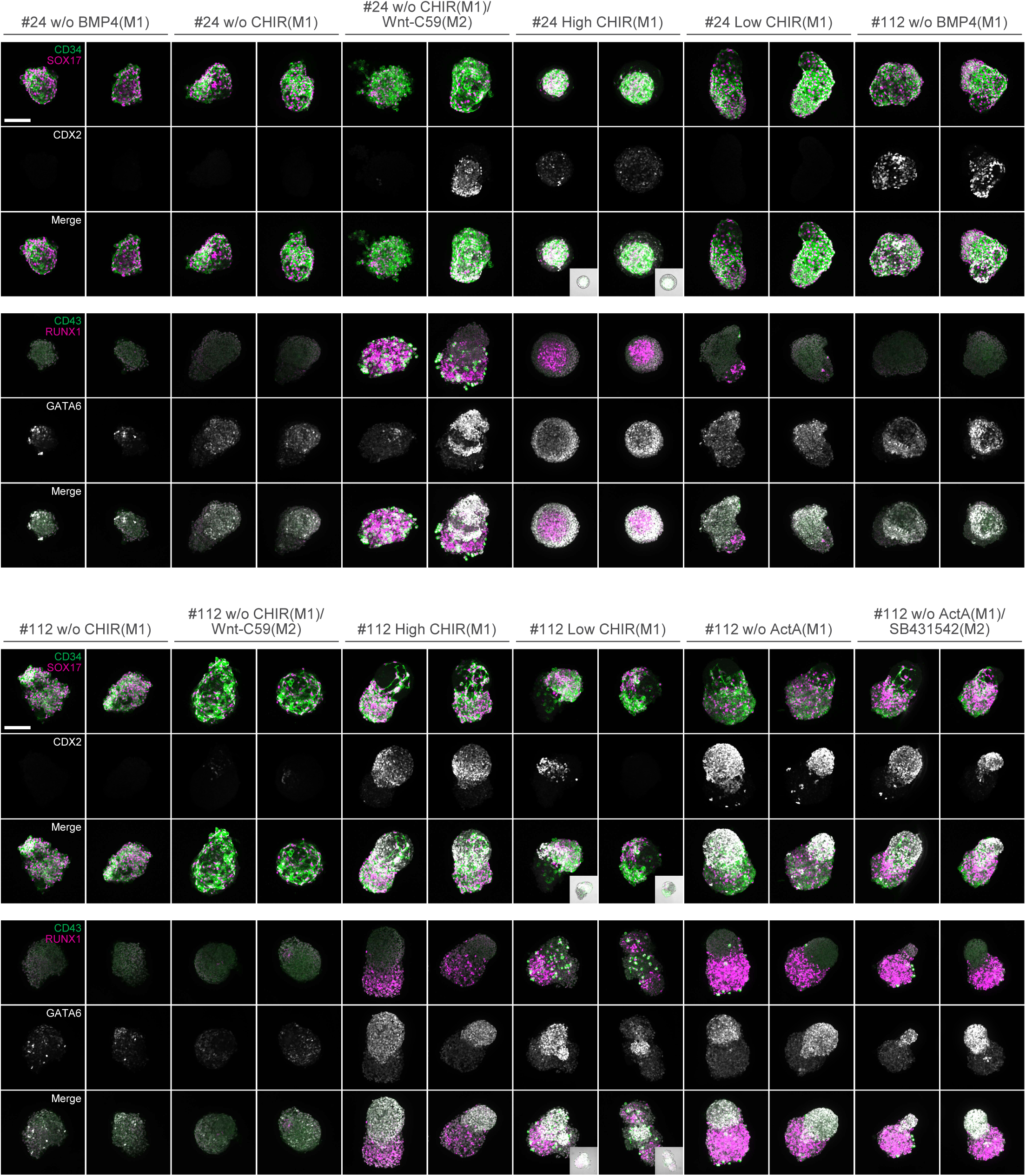
Formation of polarized EBs depending on key developmental signals. Immunofluorescence analysis of day 8 EBs cultured in the absence of key signaling molecules. Two representative maximum intensity projections of confocal microscopy images are shown for each condition. Images for control conditions (Parameter ID #24 and #112) are shown in Fig. 6b. Inset panels show merged fluorescence and bright-field images. Scale bar, 100 µm.

**Supplementary Table 1. Flow cytometry datasets for automated analysis**

**Supplementary Table 2. Process parameters, hematopoietic scores and phenotype groups for all 300 samples**

**Supplementary Table 3. Process parameters, mean hematopoietic scores, phenotype groups and mean proportions of HPC for all 246 conditions**

**Supplementary Table 4. Antibodies used for flow cytometry**

**Supplementary Table 5. Antibodies used for immunofluorescence**

